# On Casein Kinase-2 (CK2) deregulation in NSCLC: an enzyme subunit-centered approach

**DOI:** 10.1101/2023.08.04.551954

**Authors:** George V. Pérez, Li Chen, Deng Chenyi, Yin Ying, Zhao Qiang, Zhang Zhiwei, Yang Ke, Silvio E. Perea, Yasser Perera

**Affiliations:** Departamento de Bioquímica y Biología Molecular, Facultad de Ciencias Químicas, Universidad Complutense, Madrid, 28040, Spain; (G.V.P); Molecular Oncology Group, Department of Pharmaceuticals, Biomedical Research Division, Center for Genetic Engineering & Biotechnology (CIGB), Havana 10600, Cuba; (S.E.P.); Department of Pathology, Nanhua University, Hunan, China; (L.C.); (Z. Z.); The First Affiliated Hospital of the University of South China; (D.C.); (Z. Q.); China-Cuba Biotechnology Joint Innovation Center (CCBJIC), Yongzhou Development and Construction Investment Co., Ltd. (YDCI), Yangjiaqiao Street, Lengshuitan District, Yongzhou City, Hunan, 425000, China; (Y.Y.); (Y.K.); (Y.P.)

## Abstract

CK2 is considered a constitutively active protein kinase promoting/supporting several neoplastic properties and inducing a so-called non-oncogene addiction in tumor cells. Compared to the extensive body of pre-clinical research, the translational and clinical information on CK2 is still limited. The holoenzyme, composed by a tetrameric array of two catalytic (CSNK2A1 and/or CSNK2A1) and two regulatory (CSNK2B) subunits, remains to be clinically validated. Herein, we interrogated available cancer multiomics databases to unravel CK2 deregulated expression in NSCLC. We focused our analysis on individual CK2 subunits assuming subunit-specific tumor supportive roles across cancers and particularly, within two major NSCLC subtypes. Moreover, we performed meta-analysis to uncover associations between CK2 expression and patient survival, as well as further correlations analysis with components of the tumor-microenvironment. The genomic and transcriptomic data analysis was complemented by IHC evaluation of CSNK2A1, CSNK2A2 and CSNK2B subunit expression, and CK2 enzymatic activity thereof. Overall, our data suggests that epigenetic, transcriptional and post-transcriptional regulatory mechanisms rather than mutational/gene amplification events may account for differential CK2 subunits expression/activity in NSCLC. Of note, CSNK2A1 and CSNK2B mRNA up-regulation consistently determine a worse patient prognosis in LUAD and correlated with increased infiltration of MDSCs/CAFs. Importantly, we corroborated that CK2 protein subunits levels and enzymatic activity are significantly exacerbated in LUAD and LUSC, but only CSNK2A1 positively correlated with tumor size and disease stage in the analyzed patient cohort, thus supporting our transcriptomic-based correlation analysis. Finally, we concluded that CSNK2A1 alone and/or the homo-tetramer thereof may be more instrumental to support NSCLC than CSNK2A2; thus, tailored drugs against these molecular CK2 entities may achieve better therapeutic windows at least for advanced lung cancer treatment.

## 1. Introduction

CK2 is frequently deregulated in cancer and its broad tumor-supporting roles have been deemed as a “Non-Oncogene Addiction” (Ruzzene & Pinna, 2010). CK2 aberrant activity has been associated with transcriptional and post-transcriptional mechanisms, including post-translational (PTM) modifications and temporal-spatial subunits associations, as well as purported subunit specialization (Borgo *et al*., 2021). Of note, CK2 displays salient features as a constitutively active Ser/Thr-protein kinase partaking across cancer-related signaling cascades which promote aberrant cell proliferation, survival and metastasis (Trembley *et al*., 2023). The holoenzyme is composed by two catalytic CSNK2A1 or CSNK2A2 and two regulatory CSNK2B subunits, forming homo-tetramers [CSNK2A1(2)/CSNK2B(2) or CSNK2A2(2)/ CSNK2B(2)] or hetero-tetramers [CSNK2A1/CSNK2A2/CSNK2B(2)], but also building supra-molecular structures (Borgo *et al*., 2019). Finally, individual CK2 subunits may also play independent functional roles in tumor promotion (Litchfield et al., 2001; Filhol *et al*., 2015; Zonta *et al*., 2021).

Considering CK2 uncommon biochemical features (i.e., always active, variable subunit composition, substrate preferences), widespread distributed phosphorylation motive (i.e., Ser-X-X-E/D) and horizontal/vertical impact across signaling cascades and biological processes (e.g., PI3K-AKT, transcription), CK2 has generated debate on whether is a suitable oncology target, as well as about the chances of unwanted toxicity arising from its therapeutic intervention (Salvi *et al*., 2021). Such debate has also been fueled by emerging inquiries about specificity of CK2 clinical-grade inhibitor CX-4945 and CK2 essentially in cancer (Wells *et al*., 2021; Licciardello *et al*., 2021). To note, since 90’s a myriad of natural and designed chemical compounds entailing different mechanism of action have been tested (Cozza, 2017). Ultimate CK2 inhibitory compounds are SGC-CK2–1 which shows a canonical ATP-Competitive Binding and AB668 a bivalent inhibitor that binds both the ATP site and an allosteric pocket unique to CK2 (Wells *et al*., 2021; Bancet *et al*., 2022). However, thus far only two different CK2 inhibitors have reach clinical grounds, the synthetic peptide CIGB-300 and small molecule CX-4945 (Solares *et al*., 2009; Pierre *et al*., 2011). Overall, an extensive body of pre-clinical research on CK2 have been produced; however, the translational and clinical information on CK2 is still relatively limited and sparse (Strum *et al*., 2022).

Hereby, we conducted a comprehensive interrogation of available cancer multiomics databases to investigate into CK2 gene mutations and deregulated expression in clinical samples, as well as performed meta-analysis to uncover putative associations between CK2 expression and Overall Survival (OS) of Non-Small Cell Lung Cancer (NSCLC) patients. In addition, we focused on individual CK2 subunits assuming subunit-specific tumor supportive roles across cancers and particularly, within two major NSCLC subtypes i.e., Lung Adenocarcinoma (LUAD) and Lung Squamous Cell Carcinoma (LUSC). Finally, CK2 protein subunits and CK2 enzymatic activity were measured by IHC from LUAD and LUSC patients and correlated with selected clinic-pathological features. Of note, we found interesting associations between particular CK2 subunits mRNA levels and patients OS (CSNK2A1, CSNK2B), metastatic potential (CSNK2A1, CSNK2B) and tumor infiltration populations (CSNK2A1/CSNK2A2/CSNK2B), as well as among CK2 subunits levels on patient biopsies (CSNK2A1) and relevant clinic-pathological features in analyzed NSCLC patient cohort.

## 2. Materials and Methods

### 2.1. CSNK2 gene mutational burden in cancer cohorts

cBioPortal for Cancer Genomics (https://www.cbioportal.org/) was used to retrieve the mutational burden in terms of total frequency and alteration type for CSNK2A1, CSNK2A2 and CSNK2B subunits from the Pan-cancer analysis of whole genomes (ICGC/TCGA, Nature 2020) (Cerami *et al*., 2012) (Gao *et al*., 2013). Samples with mutations and copy-number amplifications (CNA) data were selected across 38 tumor types (2683 samples / 2565 patients).

cBioPortal was used to retrieve genetic alterations on CSNK2A1, CSNK2A2 and CSNK2B subunits for particular NSCLC cohorts (i.e., LUAD and LUSC). The Select Genomic Profiles were: Mutations, Putative copy-number alterations from GISTIC, and mRNA expression z-scores relative to normal samples (log RNA Seq V2 RSEM). For LUAD analysis the Lung Adenocarcinoma cohort (TCGA, PanCancer Atlas, n=566 samples) was interrogated; whereas, for LUSC analysis the lung squamous carcinoma cohort (TCGA, PanCancer Atlas, n=487 samples) were used. The CCLE were downloaded from CCLE database (https://portals.broadinstitute.org/).

Prediction of driver features based on mutational frequency, distribution and functional impact of single mutations were performed through IntOgene platform which interrogate 66 Cancer types from 221 studies (n=28,076 samples, https://www.intogen.org/search) (Martínez-Jiménez *et al*., 2020). Annotation of cancer driver features were collected from OncoKB™ - MSK’s Precision Oncology Knowledge Base (<u>OncoKB™ - MSK’s Precision Oncology Knowledge Base</u>) (Chakravarty *et al*., 2017).

### 2.2. CSNK2 genes expression in cancer

Expression levels of CSNK2 genes across pan-cancer studies were interrogated using data from the TCGA study available at The University of Alabama at Birmingham database (UALCAN) (https://ualcan.path.uab.edu) (Chandrashekar *et al*., 2017). Tumoral and non-tumoral samples were represented as red and blue, respectively. TCGA tumoral subtypes were labeled as: Bladder urothelial carcinoma (BLCA); Breast invasive carcinoma (BRCA); Cervical squamous cell carcinoma (CESC); Cholangiocarcinoma (CHOL); Colon adenocarcinoma (COAD); Esophageal carcinoma (ESCA); Glioblastoma multiforme (GBM); Head and neck squamous cell carcinoma (HNSC), Kidney chromophobe (KICH); Kidney renal clear cell carcinoma (KIRC); Kidney renal papillary cell carcinoma (KIRP); Liver hepatocellular carcinoma (LIHC); Lung adenocarcinoma (LUAD); Lung squamous cell carcinoma (LUSC); Pancreatic adenocarcinoma (PAAD); Prostate adenocarcinoma (PRAD); Pheocromocytoma and paraganglioma (PCPG); Rectal adenocarcinoma (READ); Sarcoma (SARC); Skin cutaneous melanoma (SKCM); Thyroid carcinoma (THCA); Thymoma (THYM); Stomach adenocarcinoma (STAD); Uterine corpus endometrial carcinoma (UCEC).

### 2.3. CSNK2 genes expression in NSCLC

We analyzed the expression data of CSNK2 genes using University of California Santa Cruz (UCSC) Xena (http://xena.ucsc.edu) database (Goldman *et al*., 2020). Firstly, we compared the expression of CSNK2 genes in primary tumor and normal tissue samples, using two harmonized large consortia, NCI Genomic Data Commons (GDC) and TCGA - LUAD. Then, we analyzed the expression of CSNK2 genes using the same consortia, but in this case for LUSC (GDC TCGA - LUSC). Statistical analyses were automatically computed (one-way ANOVA, p < 0.05).

### 2.4. Survival meta - analysis of CSNK2 genes in NSCLC

Survival meta-analysis of each CSNK2 gene was interrogated in LUAD or LUSC subtype explorer database (https://lce.biohpc.swmed.edu/lungcancer/index.php#about) (Cai *et al*., 2019). Output values were displayed according all curated studies available.

### 2.5. Metastasis potential of CSNK2 genes in NSCLC

Differential CSNK2 genes expression analysis in Tumor, Normal, and Metastatic tissues was carried out using TNMplot (https://tnmplot.com/analysis/) database (Bartha *et al*., 2021). Statistical methods were set automatically as Kuskal-Wallis as one-way non-parametric test.

### 2.6. Correlation of CSNK2 genes and tumor-microenvironment (TME)

TIMER2.0 (http://timer.cistrome.org) database (Li *et al*., 2020) was used to analyze correlation between CSNK2 genes expression, immune infiltrates and TME resident cells according TCGA available data. Each group of immune cells were analyzed toward purity and infiltration level. Spearman’s correlation test was the statistical method used. TCGA tumoral subtypes were labeled as: Adenocortical carcinoma (ACC); Bladder urothelial carcinoma (BLCA); Breast invasive carcinoma (BRCA); Cervical squamous cell carcinoma (CESC); Cholangiocarcinoma (CHOL); Colon adenocarcinoma (COAD); Diffuse large B-cell lymphoma (DLBC); Esophageal carcinoma (ESCA); Glioblastoma multiforme (GBM); Head and neck squamous cell carcinoma (HNSC), Kidney chromophobe (KICH); Kidney renal clear cell carcinoma (KIRC); Kidney renal papillary cell carcinoma (KIRP); Acute Myeloid Leukemia (LAML); Lower grade glioma (LGG); Liver hepatocellular carcinoma (LIHC); Lung adenocarcinoma (LUAD); Lung squamous cell carcinoma (LUSC); Mesothelioma (MESO); Ovarian serous cystadenocarcinoma (OV); Pancreatic adenocarcinoma (PAAD); Prostate adenocarcinoma (PRAD); Pheocromocytoma and paraganglioma (PCPG); Rectal adenocarcinoma (READ); Sarcoma (SARC); Skin cutaneous melanoma (SKCM); Testicular germ cell tumors (TGCT); Thyroid carcinoma (THCA); Thymoma (THYM); Stomach adenocarcinoma (STAD); Uterine corpus endometrial carcinoma (UCEC); Uterine carcinosarcoma (UCS); Uveal melanoma (UVM).

### 2.7. Essentially of CSNK2 genes

We used shinyDepMap (https://openebench.bsc.es/tool/shinydepmap) database (Shimada *et al*., 2021) to determine CSNK2 genes pan-cancer essentiality. We analyzed efficacy, selectivity and dependency scores parameters across different tested cancer cell lines. We used default algorithm of CRISPR and short-hairpin RNA (shRNA) mix ratio (CRISPR 60: shRNA 40). Then, we selected gene cluster, connectivity and correlation algorithms to predict alternative dependency score for CSNK2 genes.

### 2.8. Expression, correlation and dependency of CSNK2 genes in NSCLC

We explored the Cancer Dependency Map of CSNK2 genes throughout NSCLC cell lines using DepMap Portal (https://depmap.org/portal/) (Tsherniak *et al*., 2017). We filtered NCSLC cell lines and classified it according sub-subtype lineage. Moreover, for knockout (KO) analysis we used the integrated analysis of CRISPR (DepMap Public23Q2+Score, Chronos).

### 2.9. Immunohistochemical analysis of LUAD and LUSC specimens

A total of 116 pairs of wax blocks were prepared from 119 collected primary tumor specimens (LUAD=90; LUSC=26) or corresponding para-neoplastic tissue (>5 cm) in patients initially diagnosed at the First Affiliated Hospital of South China University since December 2021 (Hunan, China). By the time of specimen collection, the patients had not undergone radiotherapy and displayed no other systemic primary tumors except NSCLC cancer. Postoperative HE sections were diagnosed as LUAD or LUSC.

Immunohistochemical (IHC) analysis were performed by using primary antibodies against CSNK2A1, CSNK2A2, CSNK2B, NPM1/NPM1s125, and Ki67 at recommended dilutions. Immunohistochemical SP method (according to Fuzhou Maishin immunohistochemical ultrasensitive mouse/rabbit SP IHC ready-to-use kit) was performed to detect the expression of CK2A1 (Abcam 76040), CK2A2 (Invitrogen PA5-109601), CK2B (Abcam 76025), NPM1 (ab52644), NPM1s125 (ab109546) and Ki-67 (Fuzhou Maishin) proteins in human lung cancer tissues and their corresponding para-cancerous normal lung tissues. A Combined Scoring System (CIS) was implemented based on signal intensity (from non-stained=”0” to “3” points strong brown) and frequency of immunostaining (positive cells <5%=0 points up to >75%=4 points) for tumor cells. Each analyzed section was randomly selected from the upper, lower, left, right, and central five regions of magnified field (×400) for interpretation and assessment. The Combined A and B multiplied scores were as follows: Score 1-3 as (−); Score 4-5 as (+); Score 6-7 as (++); Score ≥8 as (++++). Combined scores above 1, 2, 3 and 4 correspond to negative, weak positive, positive and strong positive respectively. Statistical analysis was performed using SPSS 27.0. Chi-Squared Test and Fisher’s exact test correlation were used. A two-sided P < 0.05 value was considered to indicate statistical significance for all calculations.

## 3. Results

### 3.1. Mutational burden of CK2 subunits across cancer cohorts

CK2 deregulation has been deemed as an example of non-oncogene addition owing to its well documented supportive roles in exacerbated cell proliferation, survival and metastasis (Mandato et al., 2016). Thus, we aimed to analyze all genetic alterations on CSNK2 genes which may partake in such aberrant kinase activity by interrogating available cancer genomics databases. **Figure 1A** shows that according to cBioPortal, the percentage of genetic alterations for CSNK2A1, CSNK2A2 and CSNK2B genes in a pan-cancer cohort were 7%, 1.8% and 6%, respectively. Most of these genetic alterations comprised CNA; whereas, missense, splice site and truncating mutations were below 0.3%. Of note, such uncommon mutations neither entail oncogenic potential based on their frequency, distribution and functional impact (IntOgene) or according to annotations from a Precision Oncology Knowledge Base (OncoKB) (data not shown). Overall, genetic alterations on CSNK2A1, CSNK2A2 and CSNK2B genes tends to co-occur, both when analyzed CNA + mutations altogether (**Fig.1A**, lower panel), as well as when analyzed CNA alone (data not shown). For CSNK2A2 copy number amplifications and deletions were observed in five tumor types; whereas, CSNK2A1 and CSNK2B gene amplifications prevails in 14 out of 16 tumor localization when 3% of genetic alterations in CK2 genes were used as arbitrary threshold (**Fig.1B; SF.1B**).

**Figure 1.**
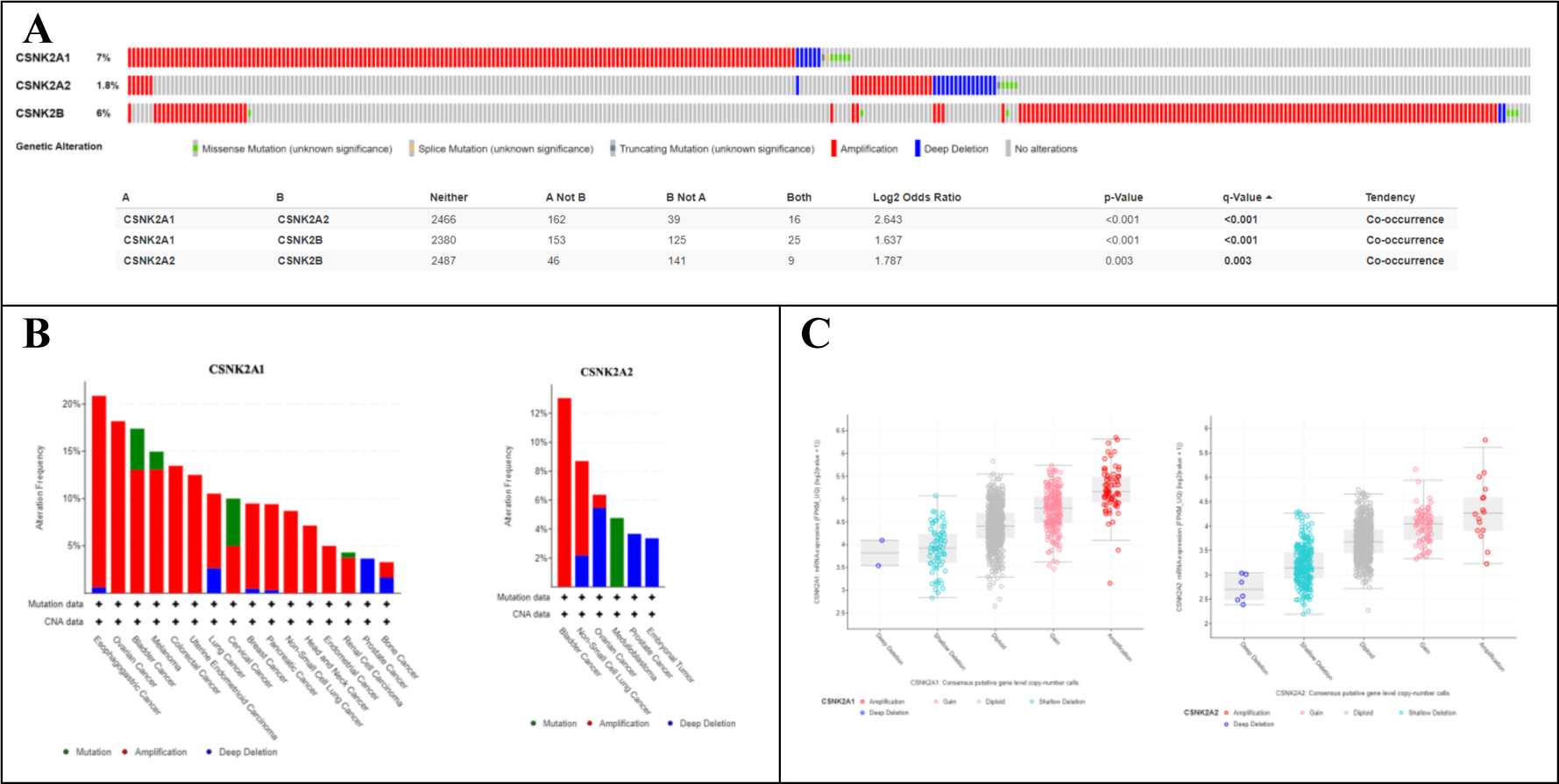
Oncoprint analysis of genetic alterations contained in pan-cancer cohorts (ICGC/TCGA, Nature 2020) for catalytic (CSNK2A1/CSNK2A2) and regulatory subunits (CSNK2B) of CK2. A) Oncoprint and co-occurrence analysis, only samples with mutation and CNA information were analyzed (92% cohort, n=2683 samples); B) Cumulative alterations on selected CK2 subunits, only those tumors with 3% or higher alteration frequency in target gene are shown; C) Correspondence among CNA (Consensus putative gene level copy-number call) and gene expression (mRNA expression, FPKM_UQ) across the database. Lung cancer (n=38) and NSCLC (n=46) data included actually comprised LUAD-US (n=35 plus other 3 unknown samples) and LUSC-US study (n=46), respectively.

On the other hand, we observed cumulative gene alteration frequencies on CK2 subunits above 10% in 15 different tumors (**SF.1A**). The higher genetic alterations in CK2 genes were reported in melanoma and bladder cancer (>30%); whereas LUAD (in the graph denoted as “lung cancer”, see **Fig.1** caption) and LUSC (in the graph denoted as “NSCLC”, see **Fig.1** caption) appeared as the fifth and sixth more altered cancer type concerning CK2 subunits (18.4%), respectively (**SF.1A**). Interestingly, in this small cohort of LUSC, all CSNK2A1 alterations were gene amplifications (4/46 cases, 9%), whereas CSNK2A2 and CSNKB harbored amplifications (3/46 cases, 7% and 2/46 cases, 4%; respectively) or deletions (1/46 cases, 2% and 2/46 cases, 4%; respectively) (**Fig.1B; SF.1B**). Overall, across this pan-cancer cohort, CNA matched with mRNA expression for the three CK2 subunits (**Fig.1C; SF.1C**). Furthermore, CSNK2A1 mRNA expression positively correlated with CSNK2A2 mRNA expression (Spearman: 0.24; p=2.10e-14), whereas an inverse correlation was found among CSNK2A1 and CSNK2B mRNA expression (Spearman: −0.15; p=2.91e-6). Finally, no correlation among CSNK2A2 mRNA and CSNK2B mRNA expression was found in this data cohort (**SF.1D-F**).

### 3.2. Genomic and transcriptomic alterations of CK2 subunits across NSCLC major subtypes

Considering that NSCLC data included in the pan-cancer cohort (ICGC/TCGA, Nature 2020) is limited, we interrogated more comprehensive datasets of lung adenocarcinoma (LUAD, n=566) and lung squamous carcinoma (LUSC, n=487). We prioritized samples containing completed data sets i.e., Mut + CNA + RNAseq (LUAD=503; LUSC=466) (**Fig.2A**); whereas, in other instances we included samples with Mutational + CNA data (LUAD=507, LUSC=469) to match **Fig.1** analysis (**Fig.2B, C**).

**Figure 2.**
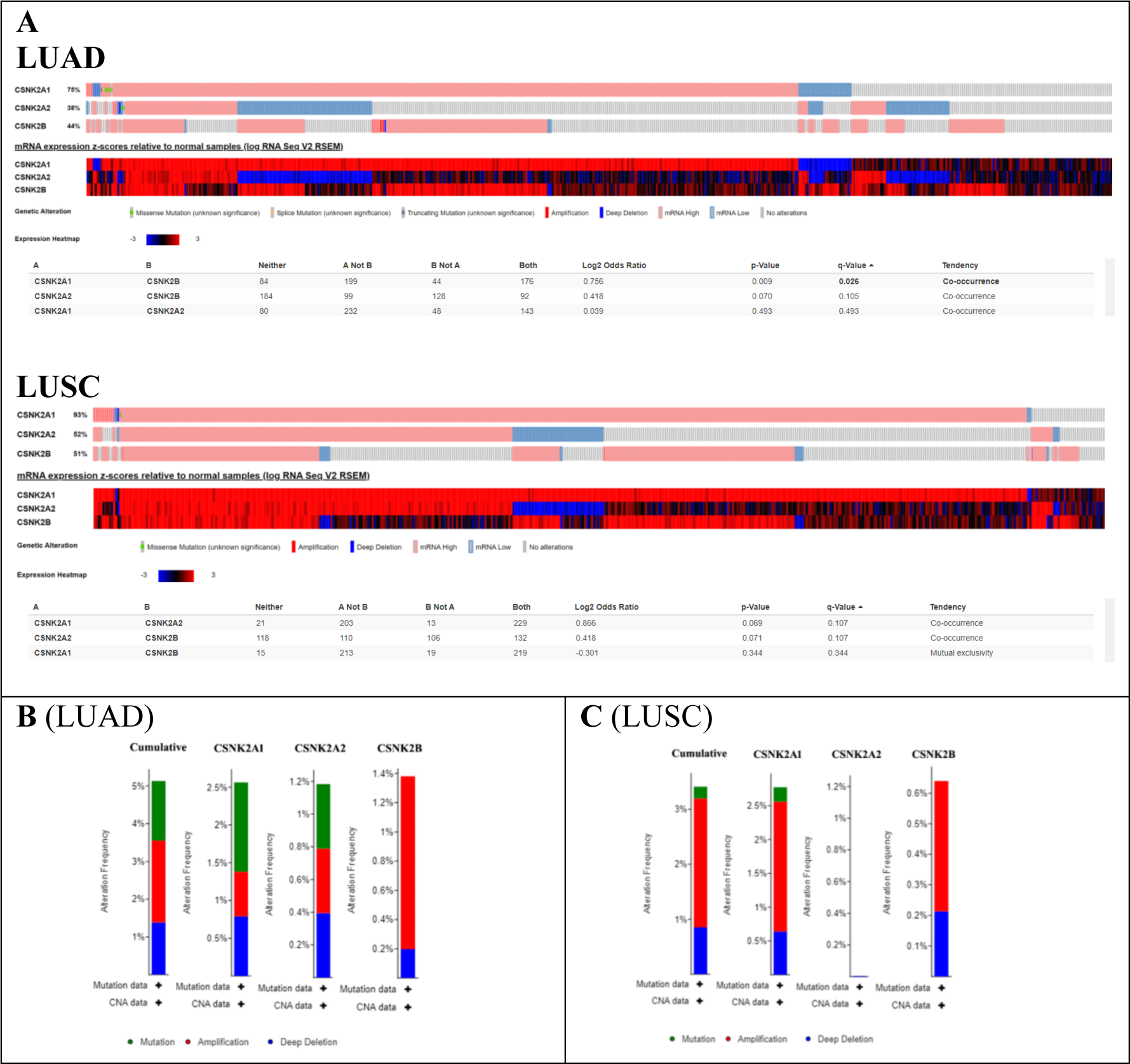
Oncoprint analysis of genetic alterations contained in LUAD and LUSC cohorts. A) Only samples reporting Mut + CNA + mRNA expression data, i.e., completed samples, were included in the analysis (LUAD=503; LUSC=466); B) LUAD samples including CNA + Mut data were included (n=507); C) LUSC samples including CNA + Mut data were included (n=469)

CSNK2A1 was target of very few mutations in both LUAD and LUSC samples (6 mutations), whereas mutations in CSNK2A2 were found only in LUAD (2 mutations). No mutations on CSNK2B regulatory gene subunit were found across 976 tested NSCLC samples (**Fig.2 B, C**). Overall, these results confirmed that mutational events on CK2 subunits CSNK2A1, CSNK2A2 and CSNK2B are rare (i.e., cumulative mutations <1.5% in tested samples) (**Fig.2 B, C**). To note, CNA events were also limited in analyzed cohorts, with amplifications and deletions only in 3-4% of samples (**Fig.2 B, C**). As in the pan-cancer cohort, such CNA roughly agreed with target gene expression (**SF.2**).

While the impact of mutations and CNA on individual CK2 gene subunits were low, corresponding mRNA over-expression abounded across analyzed LUAD and LUSC cohorts (**Fig. 2A**). Particularly, CSNK2A1 mRNA deregulation was found in more than 90% and 70% of analyzed LUSC and LUAD samples, respectively. Overall, catalytic subunits appeared more deregulated in LUSC than in LUAD samples; however, gene over-expression predominates for CSNK2A1, whereas for CSNK2A2 subunit, either high- or lower mRNA levels were found when compared to normal-matched tissues (**Fig. 2A**). Concerning CSNK2B subunit, gene over-expression is by far more common that down-regulation, with more than 40% of LUAD and LUSC samples displaying such higher mRNA levels when compared to normal-matched tissues (**Fig. 2A**).

Subsequently, we use available methylation data to seek for putative correlations between CSNK2A1-TSS200, CSNK2A2-TSS1500/1rstExon and CSNK2B-TSS1500 /1stExon /5’UTR methylation and CK2 subunits expression at mRNA levels. Negative correlations were found for CSNK2A2-TSS1500 (Spearman: −-0.15, p=8.658e-4) and CSNK2B-TSS1500/1stExon/5’UTR (Spearman: −0.14, p=1.528e-3) in LUAD (**SF.3A**). Likewise, methylation on promoter regions of CSNK2A2 and CSNK2B gene, but not CSNK2A1 gene, were also observed in LUSC samples according to signals from CSNK2A2-TSS1500 (Spearman: −0.09, p=0.0443), CSNK2A2-1rstExon (Spearman: −0.10, p=0.0368), and CSNK2B-TSS1500/1stExon/ 5’UTR (Spearman: −0.26, p=9.54e-9) (**SF.3B**).

Otherwise, to correlate CK2 subunit expression (mRNA) with previously reported upstream TFs; firstly, we analyze the protein levels of SP1, ETS1, NFKB1, STAT3 and SMAD4 TFs in LUAD and LUSC samples from the CPTAC study. SP1 was the only upregulated TFs in both neoplastic contexts (**SF.4 A;** data not shown); whereas, STAT3 was only upregulated in LUAD (data not shown). The rest of analyzed TFs were less abundant in NSCLC samples than in their corresponding non-neoplastic tissues (**Fig.3 A, B**) (**SF.4 C, E;** data not shown). To note, SP1 protein levels positively correlates with CSNK2A2 mRNA levels only in LUAD (Spearman: 0.22, p=0.02) (**SF4 B**). On the contrary, among downregulated TFs, NFKB1 protein levels negatively correlated with CSNK2A1 mRNA levels in LUAD (Spearman: −0.28, p=3.33e-3) and LUSC (Spearman: −0.45, p=3.15e-5) (**Fig.3 C, F**), as well as with CSNK2B in LUAD (Spearman: −0.41, p=9.65e-6) and LUSC tissues (Spearman: −0.23, p=0.04) (**Fig.3 E, H**). No clear correlations were found between CSNK2A2 and NFKB1 in these NSCLC subtypes (**Fig.3 D, G**). Finally, ETS1 negatively correlated with CSNK2A1 in LUSC (Spearman: −0.52, p=9.40e-7), SMAD4 displayed a negative correlation with CSNK2A2 in LUAD (Spearman: −0.31, p=9.42e-4) and STAT3 showed no correlations with CK2 subunits in NSCLC studied subtypes (**SF.4 D, F;** data not shown).

**Figure 3.**
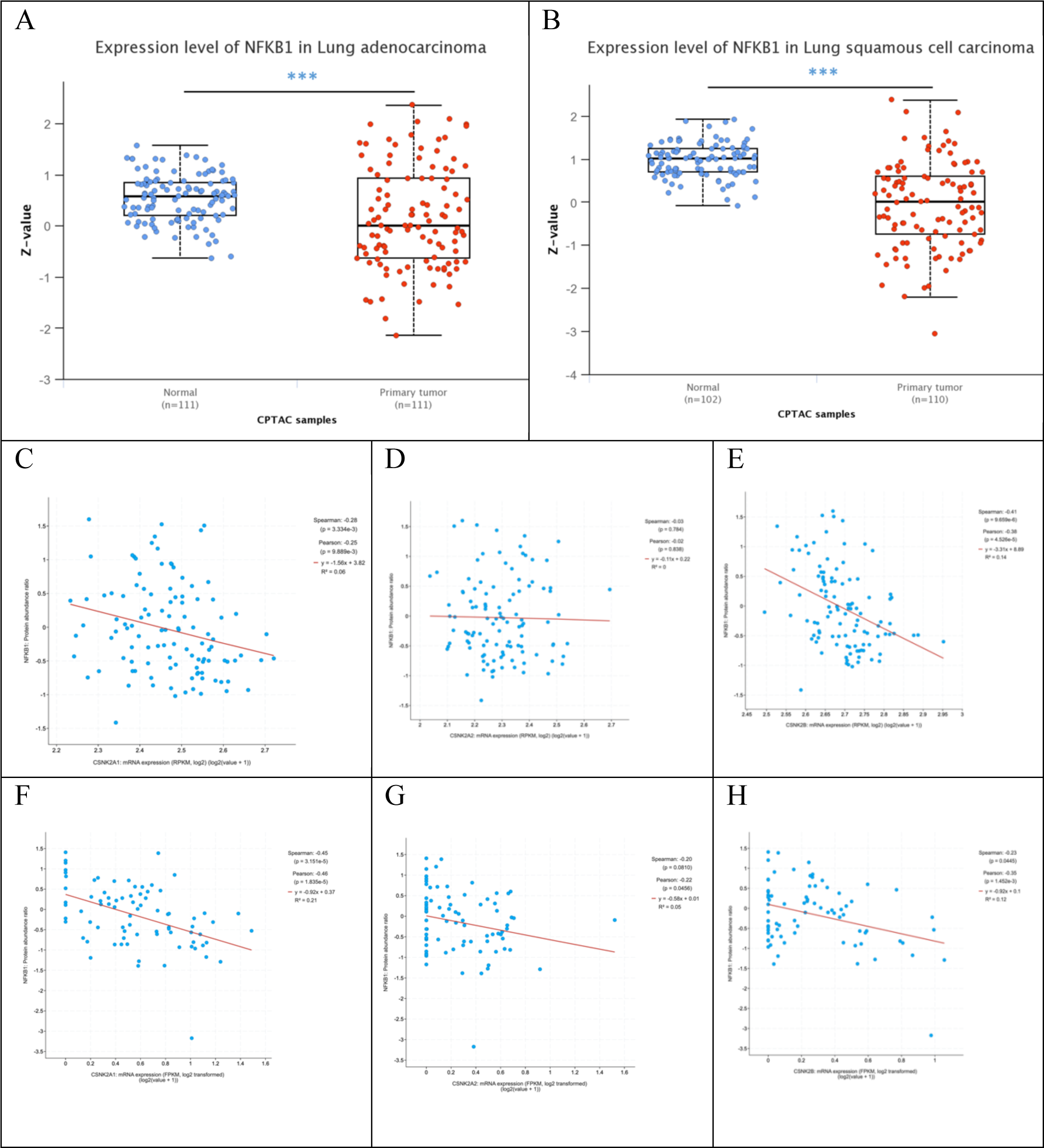
Correlations among TF NFKB1 (protein level) and CSNK2A1, CSNK2A2 and CSNK2B expression (mRNA levels) in LUAD and LUSC cohorts. Lung Adenocarcinoma and Lung Squamous cohorts (TCGA, PanCancer Atlas) containing Mass Spectrometry data from CPTAC studies (CPTAC-LUAD, n=108; CPTAC-LUSC, n=106, z-score threshold= ± 2.0) were used for TF vs mRNA correlation analysis. Expression of NFKB1 (protein) in A) LUAD and B) LUSC. Correlation between NFKB1 (protein) and CSNK2 (mRNA) expression genes in LUAD and LUSC. C, F) CSNK2A1. D, G) CSNK2A2. E, H) CSNK2B.

### 3.3. CSNK2A1 and CSNK2B over-expression decreased overall survival in LUAD

Considering that most of CSNK2A1, CSNK2A2 and CSNK2B genetic alterations can be captured by transcriptomic-based readouts, we tried to correlate CK2 subunits mRNA levels with patient survival and tumor metastasis in NSCLC clinical cohorts. Firstly, we confirmed that CK2 subunits were consistently over-expressed across tumor subtypes, including NSCLC, with the exception of CSNK2A2 in LUAD (**SF.5**). Particularly, up-regulation of CSNK2A1 was remarkably higher than CSNK2A2 in LUAD (Fold Change: 1.40 vs 0.95) and LUSC (Fold Change: 2.13 vs 1.23); whereas, CSNK2B displayed similar expression levels (LUAD: 1.35 vs LUSC:1.40) (**SF.5 C; Fig.4**). Otherwise, although data on metastatic samples is very limited as well and not stratified by tumor subtypes, it suggested that CSNK2A1 (p=1.28e-5) and CSNK2B (p=1,33e-6), but not CSNK2A2 (p=0.524), might be linked to lung tumor metastasis (**SF.6**).

**Figure 4.**
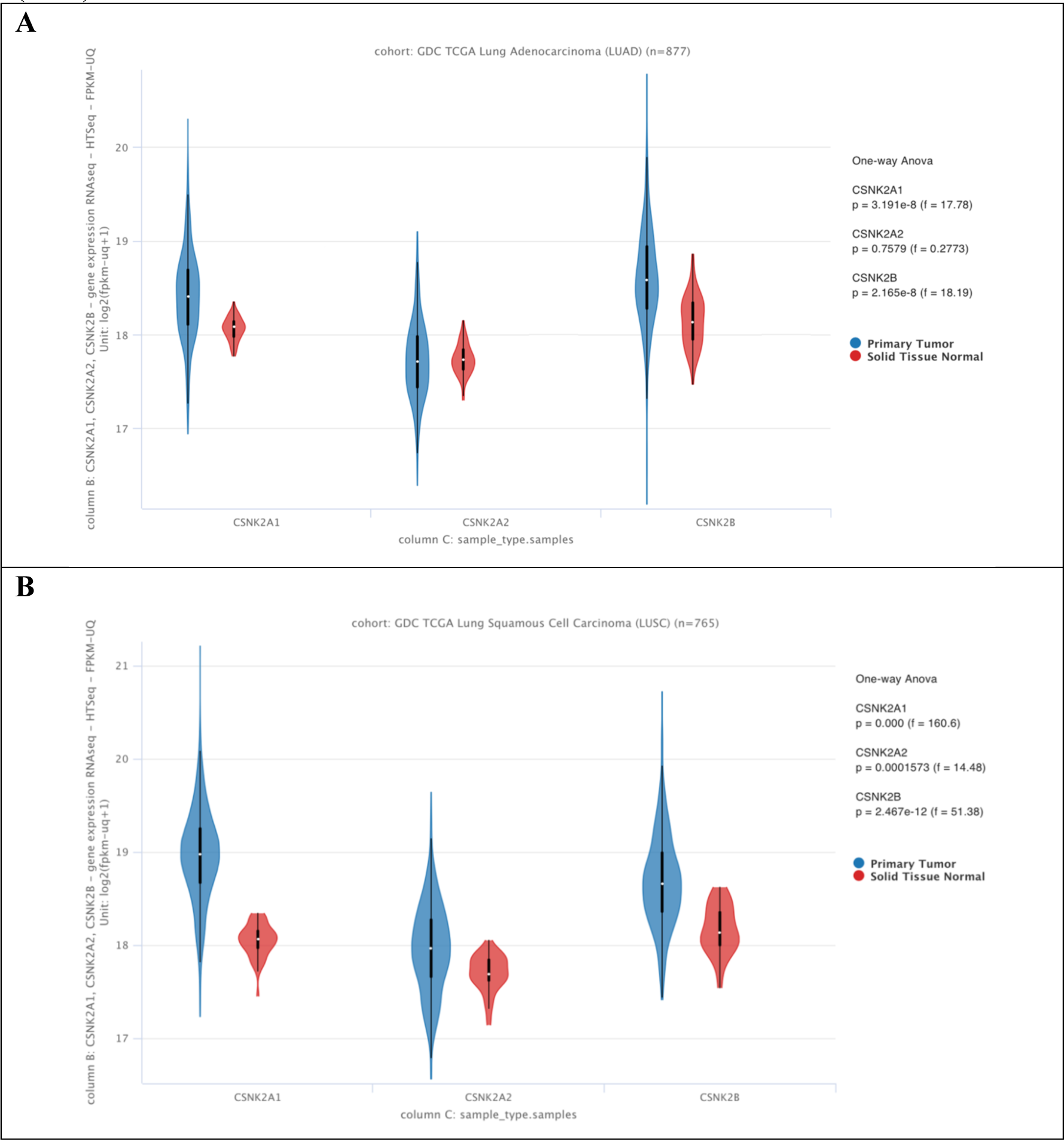
RNAseq-based expression analysis of CK2 subunits CSNK2A1, CSNK2A2 and CSNK2B in LUAD (A) an LUSC (B) patient cohorts. Normal tissue (red) and primary tumor (blue) were analyzed in some instances (LUAD). Data from GDC TCGA LUAD (877 samples) and LUSC (765 samples) were obtained from UCSC Xena database.

In line with CSNK2A1 and CSNK2B over-expression and putative metastasis association, meta-analysis of mRNA expression vs overall survival (OS) indicated that both subunits dictated worse prognosis for LUAD but not LUSC patients (CSNK2A1: LUAD HR=1.17, p<0.01; LUSC HR=1.07, p=0.72; CSNK2B: LUAD HR=1.11, p<0.01; LUSC HR=0.93, p=0.16) (**Fig.5**). Otherwise, CSNK2A2 over-expression in LUAD was associated with an increase in overall survival (HR=0.92, p=0.01) (**SF. 7)**

**Figure 5.**
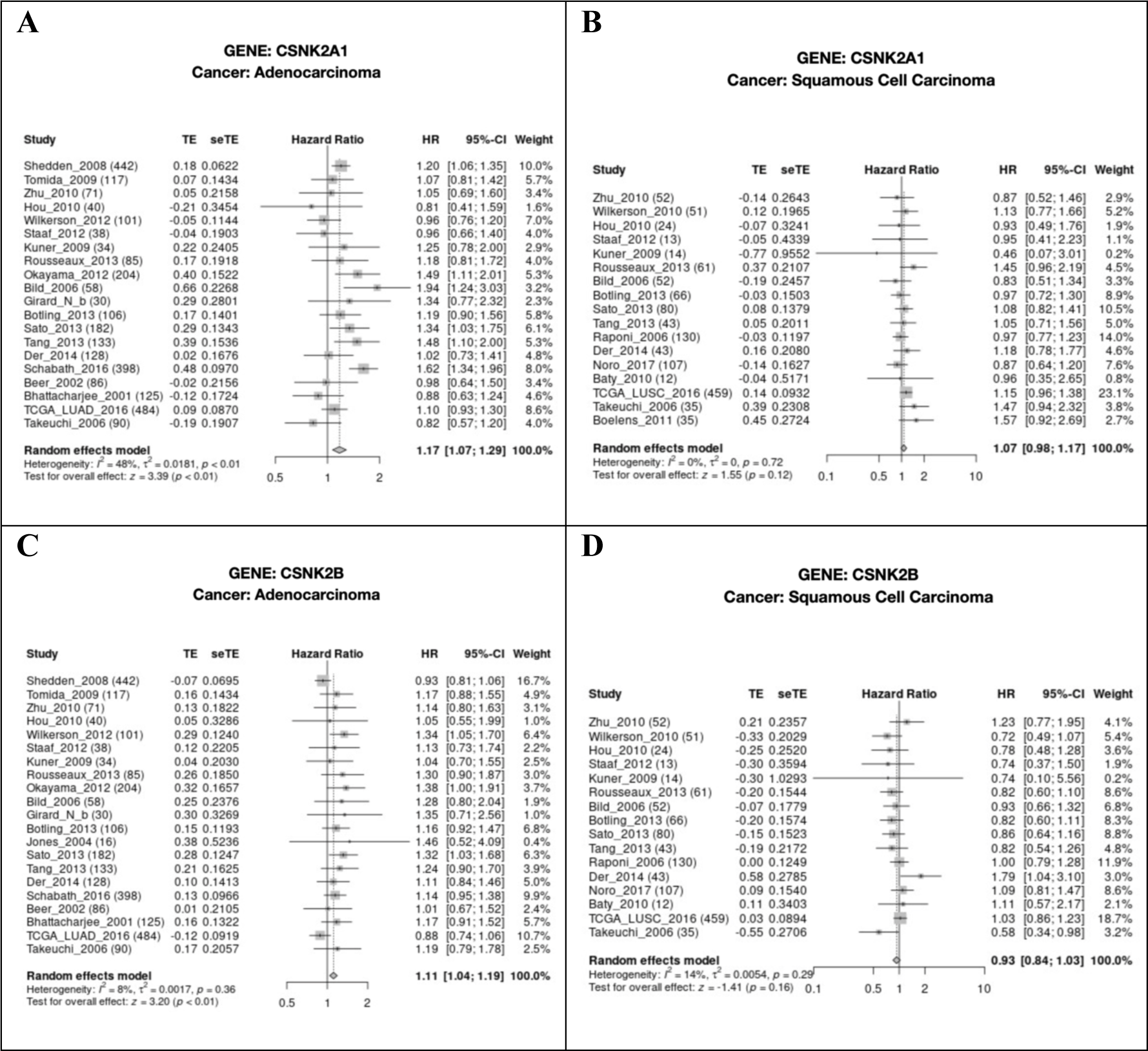
Meta-analysis of CSNK2 mRNA expression vs Overall Survival involving 20 independent data cohorts of LUAD and LUSC tumors. Hazard ratio for each CSNK2 gene was assessed independently across studies, statistics and global score was reported at left and right bottom of charts, respectively. Hazard profile of CSNK2A1 and CSNK2B in LUAD (A, C) and LUSC (B, D) respectively.

### 3.4. Over-expression of CK2 subunits correlated with increased infiltration of TME components

CK2 oncogenic potential has been linked to both tumor intrinsic and extrinsic features (Chen *et al*., 2022). By analyzing parameters related with immunosuppressive tumor infiltrating cells in clinical samples, we found striking correlations between CK2 subunits mRNA levels and tumor infiltrating cell populations. Particularly, higher CSNK2A1 mRNA levels correlated with enhanced infiltration of cancer-associated fibroblasts (CAFs) and myelo-derived suppressor cells (MDSCs) populations across several tumor subtypes (**Fig.6A**).

**Figure 6.**
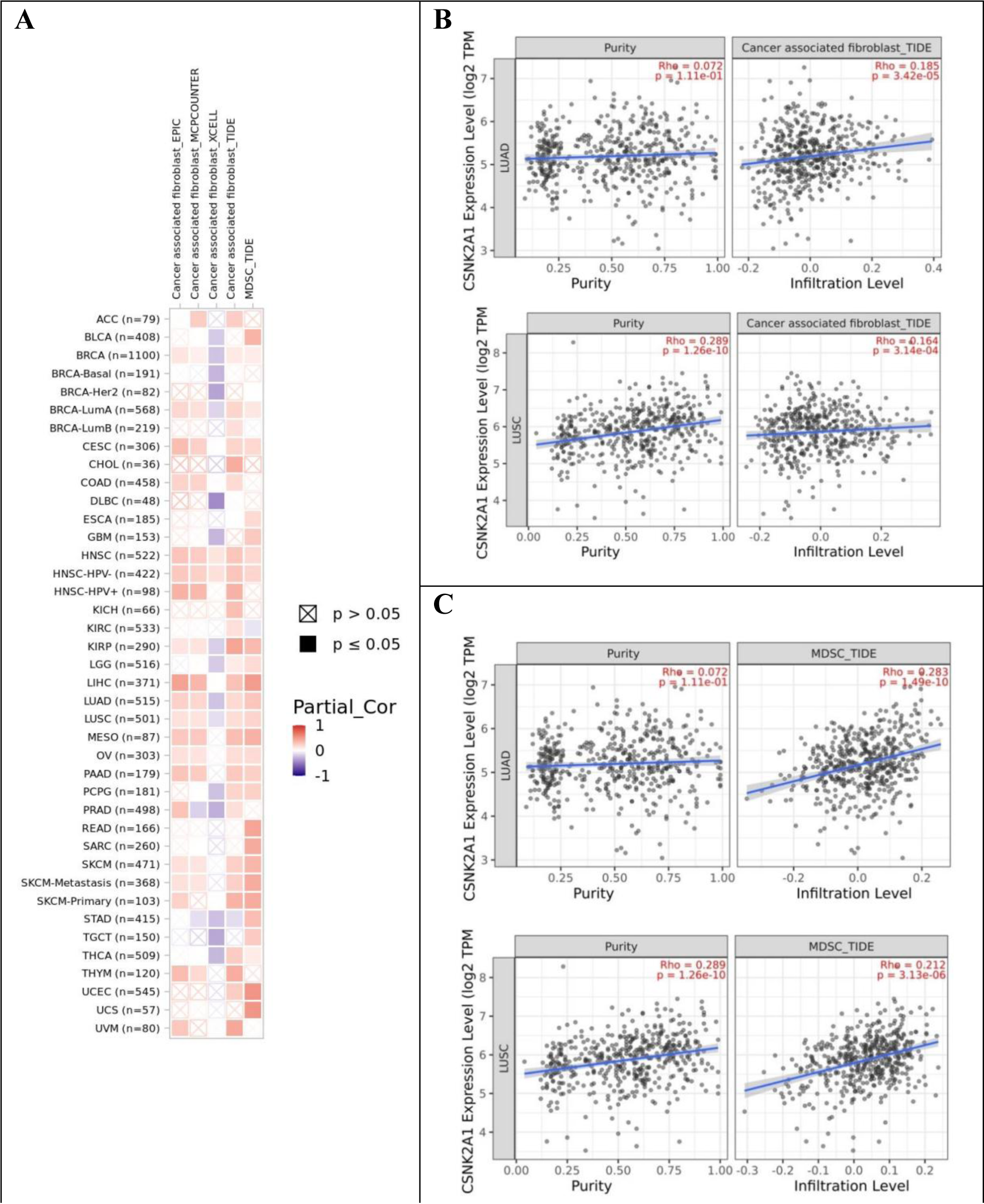
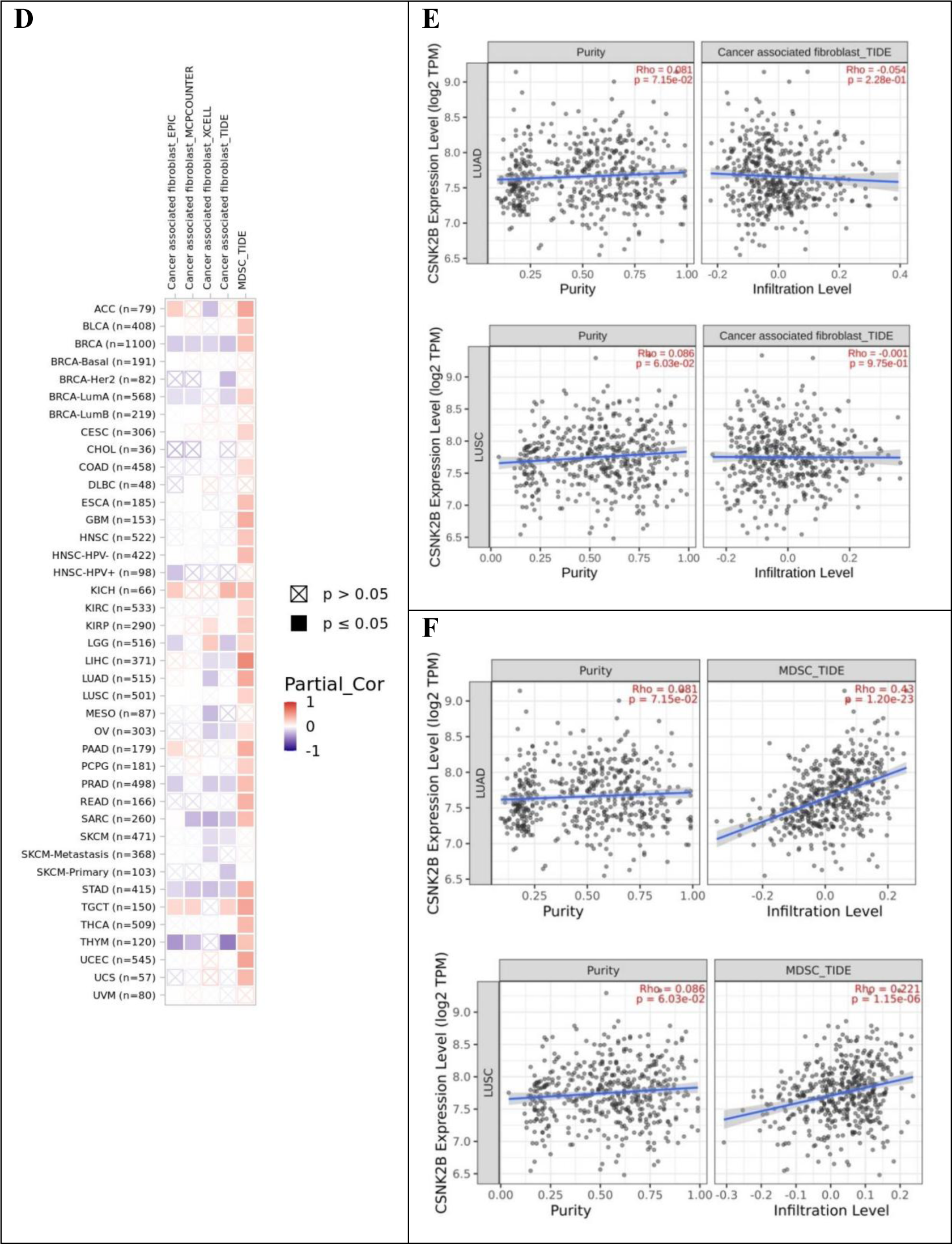
Correlation of CSNK2A1 and CSNK2B genes with selected MDSCs and CAFs in different clinical cancer cohorts. (A) Correlation of CSNK2A1 and CAFs / MDSCs throughout TCGA cancer subtypes. Detailed analysis of CAFs / MDSCs infiltration levels and CSNK2A1 expression in LUAD (B) and LUSC (C) samples. (D) Correlation of CSNK2B and CAFs / MDSCs throughout TCGA cancer subtypes. Detailed analysis of CAFs / MDSCs infiltration levels and CSNK2A1 expression in LUAD (E) and LUSC samples (F). Each analysis was normalized according the purity of the samples. TCGA subtypes are described in Materials and methods section.

Particularly, the infiltration level of these tumor-supporting cell populations correlated with CSNK2A1 expression in LUAD and LUSC subtypes (**Fig.6 B, C**). Similar correlations were observed for CSNK2A2 (**SF.8A-C**); whereas CSNK2B expression levels only correlated with infiltration level of MDSCs, reaching a remarkable correlation value (R=0.43, p=1.20e-23) in LUAD samples (**Fig.6 E, F**).

### 3.5. CSNK2B and CSNK2A1 are essential genes in cancer cell lines

Cell lines models are considered high-throughput screening platforms to validate gene essentially in cancer (Mirabelli *et al*., 2019); thus, we aimed to assess cell-dependency score for CK2 subunits using shinyDepMap database. Overall, KO of individual CK2 subunits indicated different dependency scores (DS), being CSNK2B more essential for cancer cells (−1.5<DS<-0.5), followed by CSNK2A1 (−0.4<DS<0.2) and CSNK2A2 (−0.2<DS<0.2) (**SF.9A-C**). Furthermore, the so-called “efficacy” score indicates that target CSNK2B is at least 2.5- and 3-fold more effective than targeting CSNK2A1 and CSNK2A2 genes, respectively (**Fig.7A**). In lung cancer cells CSNK2A1 and CSNK2A2 dependency scores were similar, whereas the impact on viability/proliferation of KO CSNK2B was quite significant (DS=-1.0) in 14 out of 24 tumor types (**SF.9A-C**). However, KO of CSNK2B also entailed a widespread deleterious effect in the cell panel (i.e., poor selectivity (S)=0.054); whereas, KO of CSNK2A1 was at least four times more selective (S=0.412) (**Fig.7A**).

**Figure 7.**
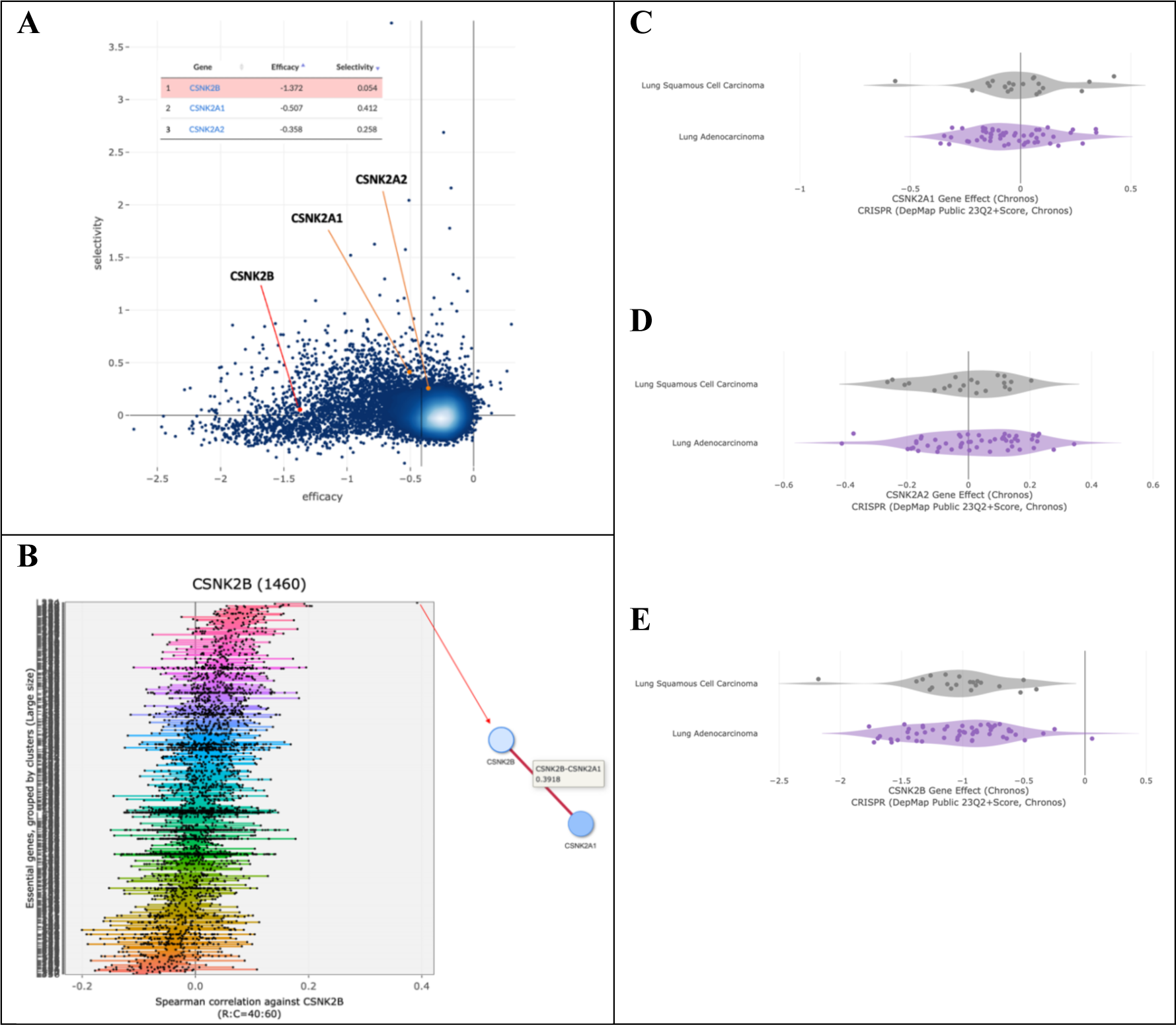
*In vitro* dependency scores for CSNK2 genes in a comprehensive cancer cell lines panel. A) Dependency map and scores for CK2 genes after KO of selected subunits across a global panel of cancer cell lines. Each subunit is organized according efficacy scores; B) Correlation for clustered genes related to CSNK2B. C-E) KO of CSNK2A1 (C), CSNK2A2 (D) and CSNK2B (E) subunits in LUAD and LUSC cells. Gene dependency effects for CNSK2A1 (A), CSNK2A2 (B) and CSNK2B (C) were retrieved from the DepMap Public 23Q2 + Score, Chronos. LUAD subtypes is represented as purple and LUSC as gray.

Subsequently, we analyzed CSNK2B clustered genes in order to find genes that may function together as complexes or pathways, thus showing similar dependency scores across cell lines (**Fig.7B**). Our findings suggested that CSNK2B and CSNK2A1 share similar dependency profiles, suggesting this particular enzyme configuration CSNK2B-CSNK2A1 may be relevant for tumor cell proliferation and survival *in vitro* (**Fig.7B, SF.9D**). Finally, by splitting KO effects across LUAD and LUSC cell lines, CRISPR of CSNK2A1 produced a slightly more deleterious effect that CSNK2A2, particularly in LUAD cells (**Fig.7 C, D**); whereas KO of CSNK2B regulatory subunit was more homogeneous and essential that its catalytic counterparts (**Fig.7E**).

### 3.6. CK2 subunits and enzymatic activity are exacerbated in LUAD and LUSC tumor specimens

To measure CK2 protein subunit levels and enzymatic activity in NSCLC tumors, we conducted IHC analysis in primary LUAD and LUSC specimens from treatment-naive patients. The intensity and frequency of CK2 subunits staining across examined tissues were variable, yet positive signals above tumor-matched para-neoplastic tissues were clearly observed (**Fig.8; SF.10**). CK2 subunits staining were detected along the nuclear and cytosolic compartments of cells; whereas, as expected Ki67 proliferation marker signal predominates in the nuclear region (**Fig.8A**).

**Figure 8.**
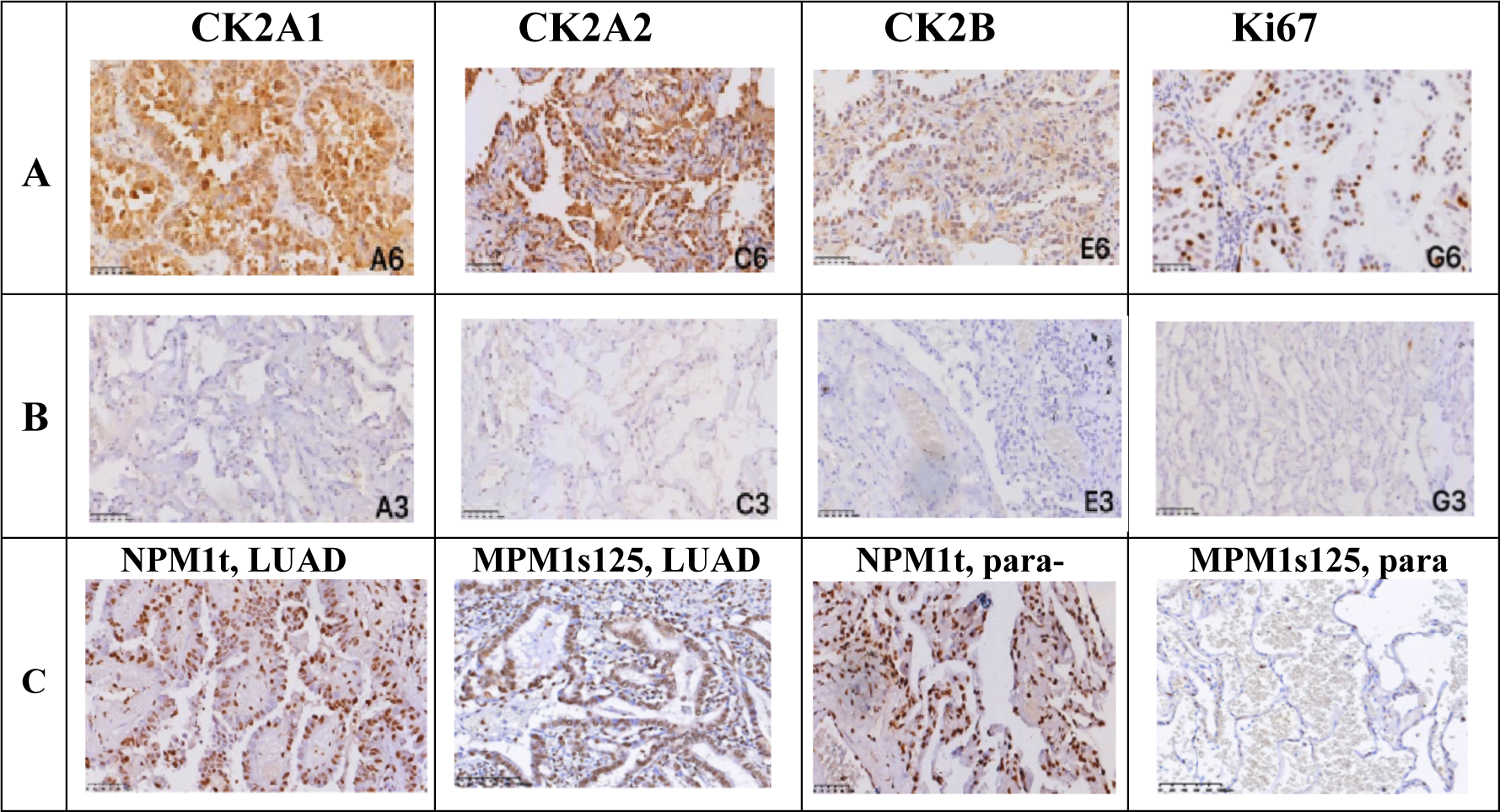
IHC detection of CK2 subunits and phosphorylation of selected CK2 substrates NPM1 in LUAD specimens and para-neoplastic tissues. Representative results for each CK2 subunits are shown in tumor (A) or para-neoplastic tissues (B); C) Total NPM1 Protein and NPM1s125 phosphorylation were measured in the same LUAD or para-neoplastic tissue of each patient to use NPM1 substrate as a proxy for CK2 in situ enzymatic activity. Representative images were taken at 200X fold magnification.

To quantify putative differences between CSNK2A1, CSNK2A2 and CSNK2B levels in tumors and corresponding para-neoplastic tissues, a Combined IHC Score (CIS), comprising signal intensity and frequency of positive cells across optical fields, was estimated for each specimen (**Table 1**).

**Table 1.** IHC detection and quantification using a Combined IHC Score (CIS) of independent CK2 subunits in primary tumor specimens of NSCLC and para-neoplastic tissues.

| Groups | Samples | CK2A1 |  |  |  | Positive rate | Z value | P value |
| --- | --- | --- | --- | --- | --- | --- | --- | --- |
|  |  | - | + | ++ | +++ |  |  |  |
| NSCLC tissue | 134 | 30 | 50 | 27 | 27 | 77.6 | -9.864 | 0.000 |
| Adjacent normal tissue | 134 | 107 | 24 | 2 | 1 | 20.1 |  |  |
| CK2A2 |  |  |  |  |  |  |  |  |
| NSCLC tissue | 134 | 16 | 55 | 41 | 21 | 88.1 | -7.248 | 0.000 |
| Adjacent normal tissue | 134 | 65 | 49 | 16 | 4 | 51.5 |  |  |
| CK2B |  |  |  |  |  |  |  |  |
| NSCLC tissue | 134 | 13 | 48 | 35 | 38 | 90.3 | -8.538 | 0.000 |
| Adjacent normal tissue | 134 | 76 | 35 | 18 | 5 | 43.3 |  |  |
| Ki-67 |  |  |  |  |  |  |  |  |
| NSCLC tissue | 134 | 58 | 32 | 20 | 24 | 56.7 | -9.722 | 0.000 |
| Adjacent normal tissue | 134 | 131 | 3 | 0 | 0 | 2.2 |  |  |

p, two-sided P < 0.05 was considered to indicate statistical significance for all calculations.

The CIS corroborated that all CK2 subunits were significantly exacerbated in NSCLC vs para-neoplastic tissues (**Table 1**). However, when we estimate the ratio (R) of Tumor(T)/Para-neoplastic (PN) positive staining (i.e., R=T/PN) as a proxy for tumor-specific association, CSNK2A1 subunit was almost two-fold more specific (R=3.9), than CSNK2B (R=2.1) and CSNK2A2 (R=1.7) counterparts. This notion was further reinforced by analyzing samples with mild-to-strong (**++/+++)** staining (**Fig.9A**). Herein, “R” values where more contrasting favoring CSNK2A1 as a tumor-specific CK2 subunit (R=18.3); whereas, CSNK2A2 and CSNK2B displayed relatively lower tumor-specificity according to our criteria (R=3.1-3.2). On the other hand, Ki67 CIS corroborate the aberrant growth of LUAD and LUSC analyzed tumors (**Table 1**). To note, CK2 catalytic subunits CSNK2A1 and CSNK2A2 were more expressed in LUSC than LUAD samples, and this pattern is more evident when considering mild-to-strong stained samples (**Fig.9B**). Subsequently, we correlate CIS for each individual CK2 subunit with selected pathophysiological features from NSCLC patients (LUAD, n=103; LUSC, n=31) (**ST1-2**). CSNK2A1 subunit expression positively correlated with tumor size and disease stage (**ST.1**); whereas, CSNK2A2 over-expression did not show correlation with analyzed parameters (**ST.2**). As illustrated in Fig.9B, both catalytic subunits were significantly more expressed in LUSC than LUAD samples (**ST1-2**).

**Figure 9.**
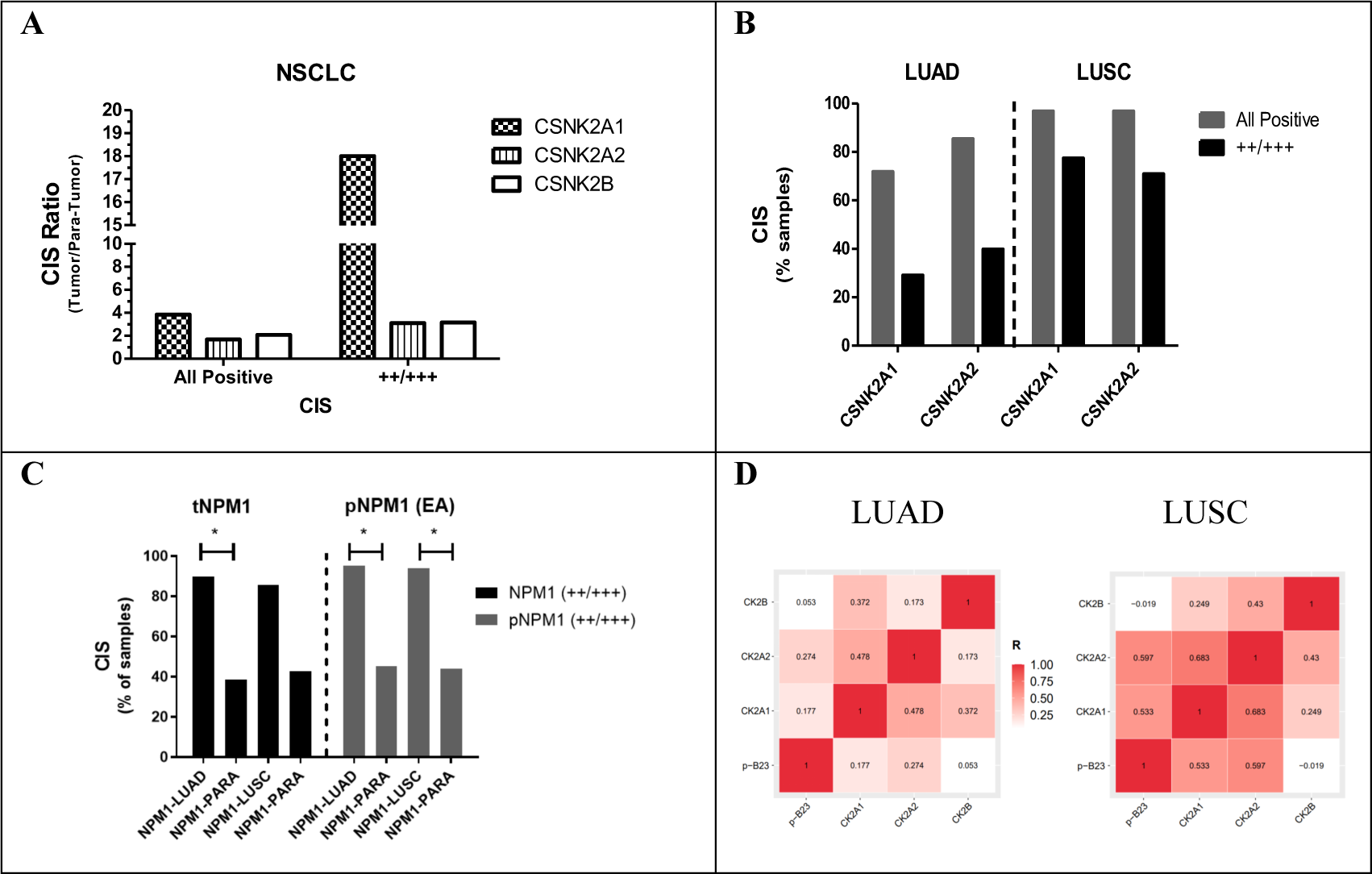
Scoring IHC results in LUAD and LUSC samples using CIS. A) Ratio (R) of Tumor(T)/Para-neoplastic (PN) positive staining (i.e., R=T/PN) as a proxy for tumor-specific association. The analysis involved scoring all positive samples or mild-to-strong intensity (++/+++).; Percent of positive samples or mild-to-strong stained samples (++/+++) in LUAD or LUSC; C) IHC results from total NPM1 protein staining (tNPM1) and NPM1s125 phosphorylated protein (pNPM1) in LUAD and LUSC specimens; D) Visualization of correlations between CSNK2A1/CSNK2A2/CSNKB protein subunit levels vs phosphorylated protein NPM1s125 levels. Statistical analysis described in ST5.

Further IHC analysis were conducted in a subset of LUAD (n=66) or LUSC samples (n=16) in order to quantify NPM1s125 phosphorylation levels as a proxy for CK2 enzymatic activity *in situ*. Both total protein levels (tNPM1), as well as phosphorylated NPM1s125 protein (pNPM1) were exacerbated in LUAD and LUSC samples compared to para-neoplastic tissues (**Fig.9C**). In both tumor types, virtually all tumor specimens (95%) showed mid-to-strong staining of pNPM1 with R>2 over para-neoplastic tissues (para-LUAD 45%; para-LUSC 44%). Interesting, pNPM1 phosphorylation levels correlated with CSNK2A2, but not with CSNK2A1, protein levels in LUAD; whereas, in LUSC pNPM1 levels correlated with both catalytic subunits CSNK2A1 and CSNK2A2 (**Fig.9D; ST5**). Otherwise, pNPM1 protein levels did not shown correlation with CSNK2B subunit content in these two major subtypes of NSCLC (**Fig.9D; ST5**). Finally, CK2 catalytic subunits at protein levels showed strong correlations among them, particularly in LUSC samples (R=0.683); whereas, CSNK2B only positively correlated with CSNK2A1 in LUAD (R=0.372) (**ST5)**

## 4. Discussion

CK2 deregulation and its purported neoplastic implications are comprehensively reviewed elsewhere (Borgo *et al*., 2021; Firnau & Brieger, 2022). Overall, enzyme biochemistry, aberrant expression, phosphorylation cascades, modulated biological processes, as well as overarching CK2 cell-intrinsic and extrinsic tumor-supportive activities, have been extensively investigated in pre-clinical models (Chua *et al*., 2017; Strum *et al*., 2022). In addition, a myriad of CK2-aiming therapeutic compounds with different inhibitory mechanisms have also been developed and tested on cell-free, *in vitro* and *in vivo* experiments (Cozza et al., 2017). However, despite a promising therapeutic potential, only two CK2 inhibitors have been investigated in clinical trials thus far (Solares *et al*., 2009; Pierre *et al*., 2011).

Of note, translational research and clinical data analysis are comparatively limited and sparse (Strum *et al*., 2022). In a pioneering study, Ortega *et al*., 2014 used Oncomine database to analyze CK2 transcript expression in seven tumor types including lung cancers. In this clinical indication CSNK2A1, CSNK2A2 and CSNKB mRNA transcripts were found over-expressed in 14/14, 2/2 and 3/3 significant unique analysis. Subsequently, Kaplan-Meier analysis only showed significant correlations between CSNK2B over-expression and lower OS, as well as the somehow counter-intuitive finding that CSNK2A2 over-expression correlates with an improved survival in LUAD (Ortega *et al*., 2014). A subsequent analysis involving another 16 tumor localizations illustrated some cancer-type specific up- or down-regulation of CK2 transcripts, as well as direct and inverse correlations among CK2 transcripts and OS (Chua *et al*., 2017). However, the prevailing picture is an increase in CK2 transcript, protein and activity levels across different cancer types which can be inhibited by cancer therapy (Strum *et al*., 2022). At the same time is anticipated that the efficacy of CK2 inhibition can be context-dependent; thus, clinical target selection must be based on best available pre-clinical, translational and clinical data (Strum *et al*., 2022). Finally, emerging evidences on CK2 subunit-specific contributions to certain tumor phenotypes stress the necessity to individually analyze their regulatory patterns and functional implications within selected tumor types (Filhol *et al*., 2015; Liu *et al*., 2016; Zonta *et al*., 2021).

Despite availability of clinical multiomics databases, a comprehensive analysis spanning mutational burden, gene/protein/activity levels deregulation and pathophysiological features correlations for individual CK2 subunits in cancer is still missing. In parallel, interrogation of well-characterized cell panels to identify purported CK2 subunit-specific cancer dependencies may complement such inquires. Altogether, these high-throughput analysis shall provide fresh insights to a CK2 community which still debates on whether CK2 is an actionable and safe oncology target to intervene (Salvi *et al*., 2021; Licciardello *et al*., 2021).

Interestingly, we found that mutational events including single base insertion, substitution and/or deletions within CK2 gene subunits seldom occur and do not entail an oncogenic potential. These findings, go along with the CK2 cooperating roles within the so called “non-oncogene addiction” (Ruzzene & Pinna 2010). Furthermore, CNA on CK2 genes are also infrequent and do not explain the broad mRNA deregulated expression observed in LUAD and LUSC patients cohorts. To investigate into other regulatory events, we selected two independent studies for each NSCLC subtype on cBioPortal and inquired for correlations between gene methylation and cognate transcription factor vs mRNA levels for each independent CK2 subunit.

The promoters of the three CK2 subunits display features of housekeeping genes including a CpG island around exon 1 which could be methylated (Ackermann *et al*., 2010). Accordingly, promoter/1stExon methylation correlated with lower CSNK2A2 and CSNK2B mRNA levels in analyzed LUSC and LUAD cohorts. Notably, the frequency of samples showing CSNK2A1 mRNA over-expression almost duplicates those displaying CSNK2A2 or CSNK2B high mRNA levels in these patient cohorts. Therefore, the lower frequency and expression levels of CSNK2A2 and CSNK2B mRNA compared with CSNK2A1 may be at least partially explained by a differential promoter methylation on CSNK2A2 and CSNK2B subunits across LUAD and LUSC analyzed samples.

Subsequently, we seek for putative correlations between TFs linked to CK2 regulation in cancer and the expression levels of CK2 subunits in LUAD and LUSC (Borgo et al., 2021). Sp1 and Ets1 were reported to bind promoters of the three CK2 subunits stimulating their transcription, while NFKB1 activates only CSNK2A1 promoter (Krehan *et al*., 2000; Krehan *et al*., 2001; Ackermann *et al*., 2005). In some instances, Ets1 acts as an oncoprotein linked to cell differentiation and invasion (Dittmer, 2003). Otherwise, compelling experimental findings evidenced the transcriptional regulation of CSNK2A1 by the STAT3 is some tumors (Kalathur et al., 2015). Finally, SMAD4 functions as a CK2 transcriptional repressor and is deleted in several tumor types including LUAD and LUSC (Zhao *et al*., 2018).

Our expression/correlation analysis indicate that NFKB1 (i.e., DNA binding subunit of NF-kappa-B protein complex or p50) is the most consistently associated TF (protein level) vs CK2 subunits expression (mRNA level) in LUAD and LUSC (inverse correlations). NFKB1 is considered a tumor suppressor gene since p50:p50 homo-dimers decrease tumor-associated inflammatory responses and anti-apoptotic signals (Elsharkawy *et al*., 2010; Schmitt *et al*., 2011). As a matter of fact, p50:p50 homo-dimers are generally transcriptional repressor by out-competing p50(NFKB1):p65(RelA) heterodimers or p65:p65 homodimers in their binding to cognate response elements in promoters (Concetti & Wilson, 2018). Such “productive” NFKB dimers mediate the transcription of several target genes including CSNK2A1 (Kheran *et al*., 2000; Giridharan & Srinivasan, 2018). Interestingly, NFKB1^−/−^ double *knockout* also favoured lung tumor progression *in vivo* by a mechanism which is independent of NFKB functions (Sun *et al*., 2016). Altogether, we believe that down-regulation of NFKB1 might unleash a transcription-based pro-inflammatory program along with CSNK2A1/CSNK2B up-regulation in both NSCLC subtypes, therefore contributing to poor patient prognosis. Interestingly, no correlations were seen between NFKB1 protein levels and CSNK2A2 mRNA expression, which match previous findings indicating CSNK2A2 gene transcription is not regulated by NFKB1 (Ackermann et al., 2005).

On the other hand, despite SP1 has been involved in coordinated transcription of all CK2 subunits and it was found significantly upregulated in LUAD and LUSC, SP1 protein levels only positively correlates with CSNK2A2 mRNA expression in LUAD. Overall, transcription-level regulation of CK2 subunits in NSCLC seems complex and multilayered with three of the five analyzed TFs showing differential subunit expression regulation within and across tumor subtypes (i.e., SP1, ETS1, SMAD4); whereas, only NFKB1 showed a consistent regulation over CSNK2A1/CSNK2B subunits in LUAD and LUSC samples.

Another emerging layer on CK2 regulation reported in clinical cancer databases is post-transcriptional control. At least three miRNAs may regulate CK2 expression in lung cancer (i.e., miR-125b, miR-532-5p, miR-217 (http://www.mirdb.org/) (Liu & Wang, 2019; Yuhao & Wang, 2020). Interestingly, a preliminary inspection indicated that most of significantly deregulated miRNAs in LUAD and LUSC tumors target CSNK2A1. Altogether, CK2 subunit-specific epigenetic gene silencing (CSNK2A2, CSK2B), TFs deregulation (e.g., NFKB1 on CSNK2A2/CSK2B) and miRNAs-based suppression (e.g., on CSNK2A1), could explain the observed differential subunit expression across LUSC and LUAD patient cohorts. However, it shall be noted that extensive post-translational modifications on CK2 subunits and cognate TFs like SP1, EST1, NFKB1, as well as protein-protein interactions add further complexity to the multilayered regulation of CK2 in cancer cells (Borgo *et al*., 2021).

Interestingly, a comprehensive meta-analysis involving 20 LUAD and 17 LUSC clinical studies, corroborated that CSNK2A1 and CSNK2B subunit over-expression entails a worse prognosis in LUAD, whereas on the contrary, CSNK2A2 over-expression associated with a better outcome. The same trend for CSNK2B and CSNK2A2 was previously found in a microarray-based study (Ortega *et al*., 2014). Overall, we found that CSNK2A1 and CSNK2B mRNA levels were exacerbated in LUAD and LUSC primary samples, as well as in a small number of available metastatic NSCLC clinical samples. In addition, we found striking direct association between the mRNA levels of CSNK2A1 and CSNK2B subunits and MDSCs/CAFs cells tumor infiltration. CK2 aberrant expression has been previously associated with poor anti-tumor immunity (Chen *et al*., 2022).

Numerous studies highlighted the importance of MDSCs/CAFs infiltration and expansion across oncogenic niches, which are thus considered targets for TME-based therapy (Yang *et al*., 2020; Wong *et al*., 2022; Shintani *et al*., 2023). These heterogeneous groups of immature myeloid cells and transformed fibroblasts display an array of cancer-supportive functions (Chen *et al*., 2021; Sheida *et al*., 2022). The accumulation of MDSCs/CAFs mainly depends on secreted molecules from tumor and transformed TME-resident cells (Wu *et al*., 2021; Glabman *et al*., 2022; Umansky *et al*., 2016; Grabilovich *et al*., 2009). Of note, many of these MDSCs/CAFs-stimulating mediators involve directly or indirectly CK2 substrates (Husain *et al*., 2021).

CK2 pharmacologic and genetic inhibition have been mostly performed *in vitro*, with few studies addressing the role of individual CK2 subunits *in vivo* (Strum *et al*., 2022). Herein, we analyzed the essentially of each CK2 subunit after KO experiments across a comprehensive cancer cell panel. KO of CSNK2B, followed by CSNK2A1 and CSNK2A2 subunits, proved essential for *in vitro* cancer cell proliferation/survival. Since KO of CSNK2B shall fully abrogate Class-III substrate phosphorylation, independent subunit functions, as well as may entail catalytic subunit instability, these results were somehow expected. Otherwise, KO of individual catalytic subunits may be attenuated by some functional overlap, particularly on *in vitro* experimental settings (Gyenis and Litchfield, 2008; Nuñez de Villavicencio-Diaz *et al*., 2017).

In contrast to abundance of clinical genomic/transcriptomic data on CK2, protein information concerning CK2 subunits and/or enzymatic activity in human lung specimens is limited. According to a recent systematic review by Strum et al. 2022 only four studies measuring CK2 levels/activity in LUAD and/or LUSC patient samples have been reported (Daya-Makin et al., 1994; Yaylim and Isbir, 2002; Liu et al., 2016; Xie et al., 2018). Liu et al. (2016) analyzed by IHC CSNK2A2 subunit expression levels in 160 NSCLC patients and found significant up-regulation of this catalytic subunit in both LUAD and LUSC specimens, marginally reducing the OS of the patients. On the other hand, the Human Protein Atlas (HPA) database contains 6 LUAD and 6 LUSC specimens (https://www.proteinatlas.org/). Medium to moderate staining for CSNK2A1 were reported in 5/6 LUAD and 3/6 LUSC specimens; whereas, all 12 specimens showed medium to high expression of CSNK2B. No IHC data was reported for CSNK2A2 in the HPA. In our work we found increased protein levels for the three CK2 subunits in a NSCLC patient cohort comprising LUAD and LUSC specimens, but importantly we demonstrate for the first time that such CK2 subunit up-regulation correlates with an increased enzymatic activity in situ, by using an endogenous CK2 substrate as proxy for such enzymatic activity.

Overall, CSNK2A1 over-expression correlates with tumor stage, size and proliferation score of NSCLC tumors. Such positive correlations agree with results from the meta-analysis where CSNK2A1 over-expression, decreased the OS of LUAD patients. Surprisingly, CSNK2A1 expression seems by far more tumor-specific than CSNK2A2 and CSNK2B in both LUAD and LUSC specimens. These findings go along with *in vitro* estimated global dependency scores for individual CK2 subunits. In such experiments, selectivity scores for CSNK2A1 were 1.6- and 7.6-fold higher than for CSNK2A2 and CSNK2B, respectively. Furthermore, CSNK2B and CSNK2A1 share similar dependency profiles, suggesting that a particular enzyme configuration composed by CSNK2A1-CSNK2B may be more relevant for tumor cell proliferation and survival *in vitro*.

All in all, our data suggests than epigenetic, transcriptional and post-transcriptional regulatory mechanisms rather than mutational and CNA events may account for differential and aberrant CK2 subunits expression/activity in NSCLC. Furthermore, that CSNK2A1 alone and/or composing a [CSNK2A1]_2_-[CSNK2B]_2_ homo-tetramer may be more instrumental to support NSCLC than CSNK2A2, particularly in LUAD tumors. Tailored drugs against CSNK2A1 or the homo-tetramer thereof, may also decrease the risks of unwanted toxicity and achieve better therapeutic windows since CSNK2A2 catalytic subunit was also abundantly expressed in normal lung cells and thus may be related to more physiologic functions therein. Finally, *in vitro* dependency scores for CSNK2 genes also hint that therapeutic strategies aiming at CK2 inhibition shall be focused in restoring back “physiological” CK2 signaling levels, rather than to fully abolish its enzymatic activity which might lead to narrow therapeutic windows in the path to the definitive validation of CK2 as oncology target in lung cancers.

