## Supplementary Material for "On Casein Kinase-2 (CK2) deregulation in NSCLC: an enzyme subunit-centered approach"

### Supplementary Materials:

**SF.1 Mutational and CNA burden for CSNK2A1, CSNK2A2 and CSNK2B included in Pan-cancer cohort (ICGC/TCGA, Nature 2020).** A) Cumulative burden in all CK2 subunits, only those tumors with 10% or higher cumulative alteration frequency in the target gene is shown; B) burden on CSNK2B, only those tumors with 3% or higher alteration frequency in target gene is shown; C) Correlation among CNA and mRNA gene expression for CSNK2B; D-F) mRNA gene expression correlations among CK2 subunits in the study cohort.

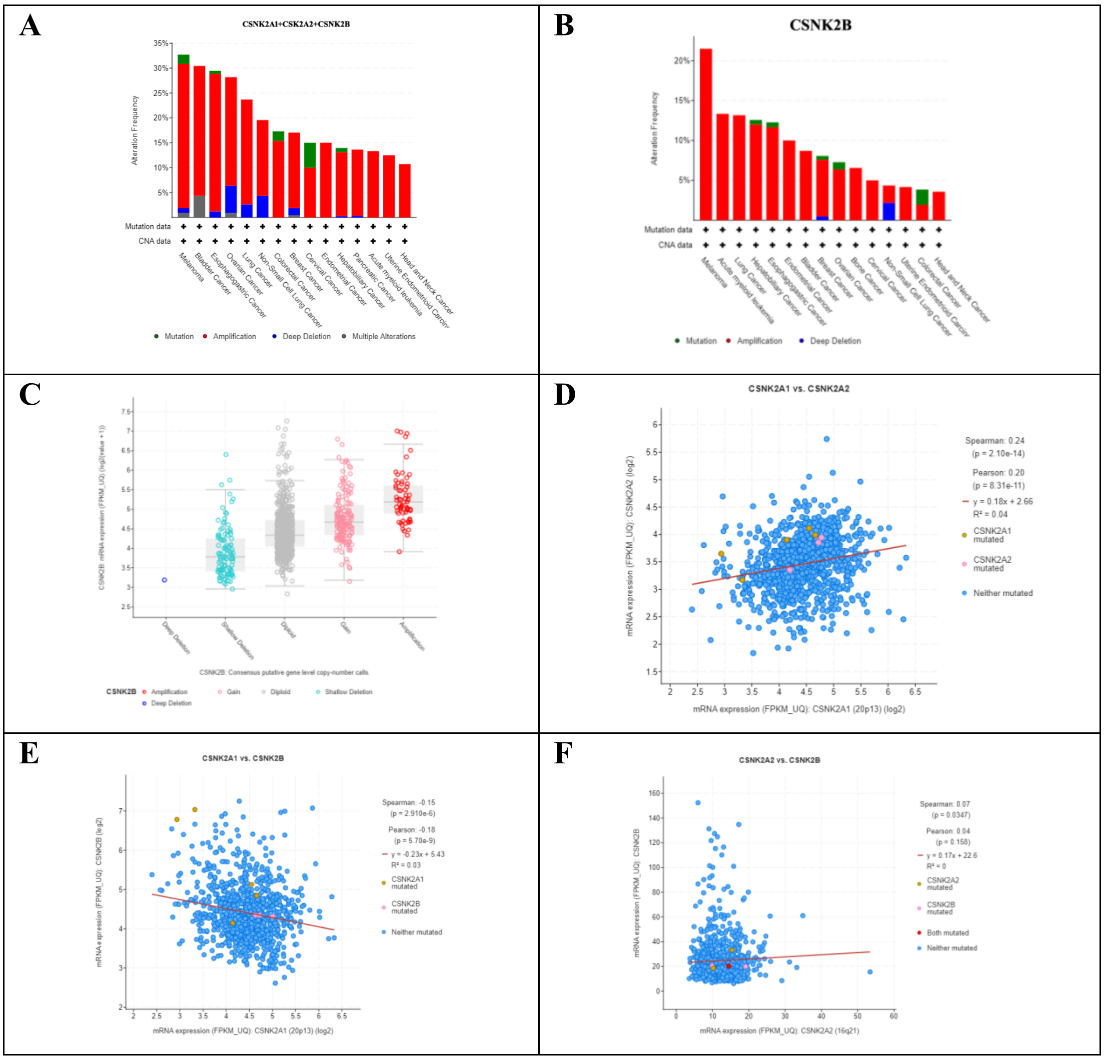

**SF.2 Association between mRNA expression and CNA in two major NSCLC subtypes.** Lung adenocarcinoma (LUAD, n=503) and Squamous Carcinoma (LUSC, n=466) cohorts with complete datasets were used in the analysis.

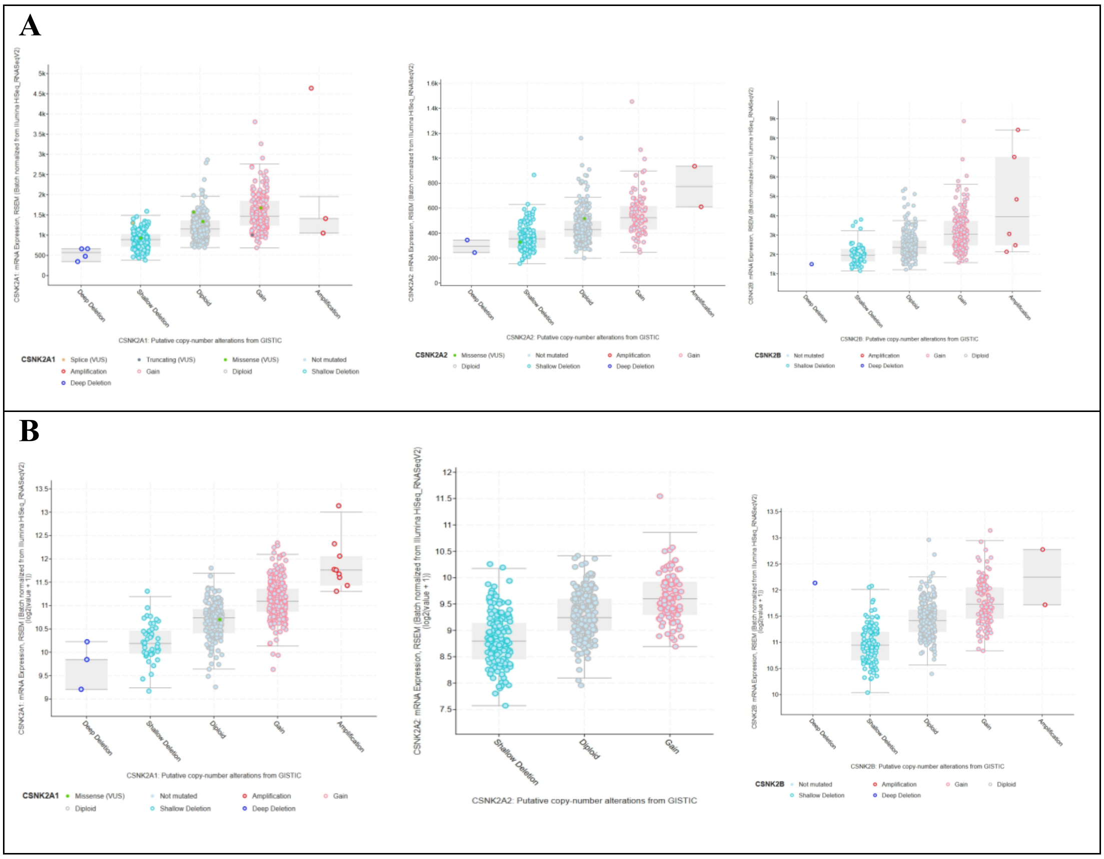

**SF.3 Correlations between Promoter/1stExon/5'UTR methylation and mRNA expression for CK2 genes in A) LUAD and B) LUSC samples.** Methylation data were collected from HM27&HM450-Merge using probes for TSS/1stExon/5'UTR for genes CSNK2A1 (TSS200), CSNK2A2 (TSS1500/1rstExon) and CSNK2B (TSS1500/1stExon/5'UTR).

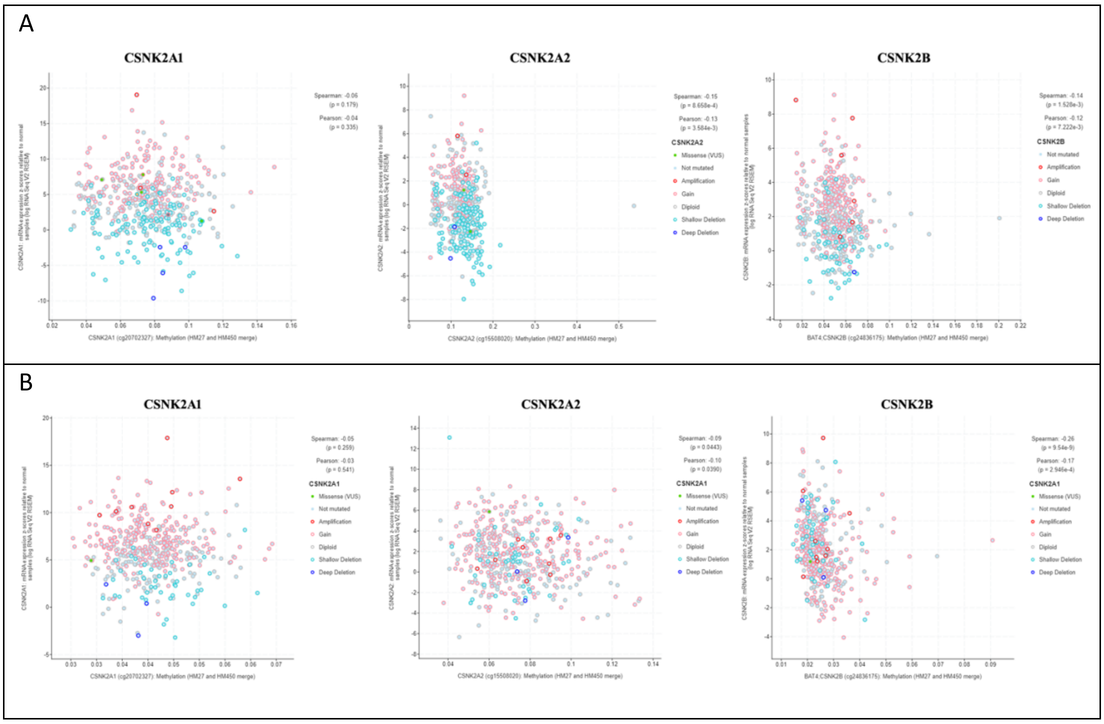

**SF.4 Correlations among upstream TFs (protein level) and CSNK2A1, CSNK2A2 and CSNK2B expression (mRNA levels) in LUAD and LUSC cohorts.** Lung Adenocarcinoma and Lung Squamous cohorts (TCGA, PanCancer Atlas) containing Mass Spectrometry data from CPTAC studies (CPTAC-LUAD, n=108; CPTAC-LUSC, n=106, z-score threshold=±2.0) were used for TFs vs mRNA expression correlation analysis. Only biologically significative correlation values between TFs and CK2 subunits are represented. A) Expression of SP1 (protein) in LUAD. B) Correlation between SP1 (protein) and CSNK2A2 (mRNA) expression in LUAD. C) Expression of ETS1 (protein) in LUSC. D) Correlation between ETS1 (protein) and CSNK2A1 (mRNA) expression in LUSC. B) Correlation between SP1 (protein) and CSNK2A2 (mRNA) expression in LUAD. C) Expression of SMAD4 (protein) in LUAD. D) Correlation between SMAD4 (protein) and CSNK2A2 (mRNA) expression in LUAD.

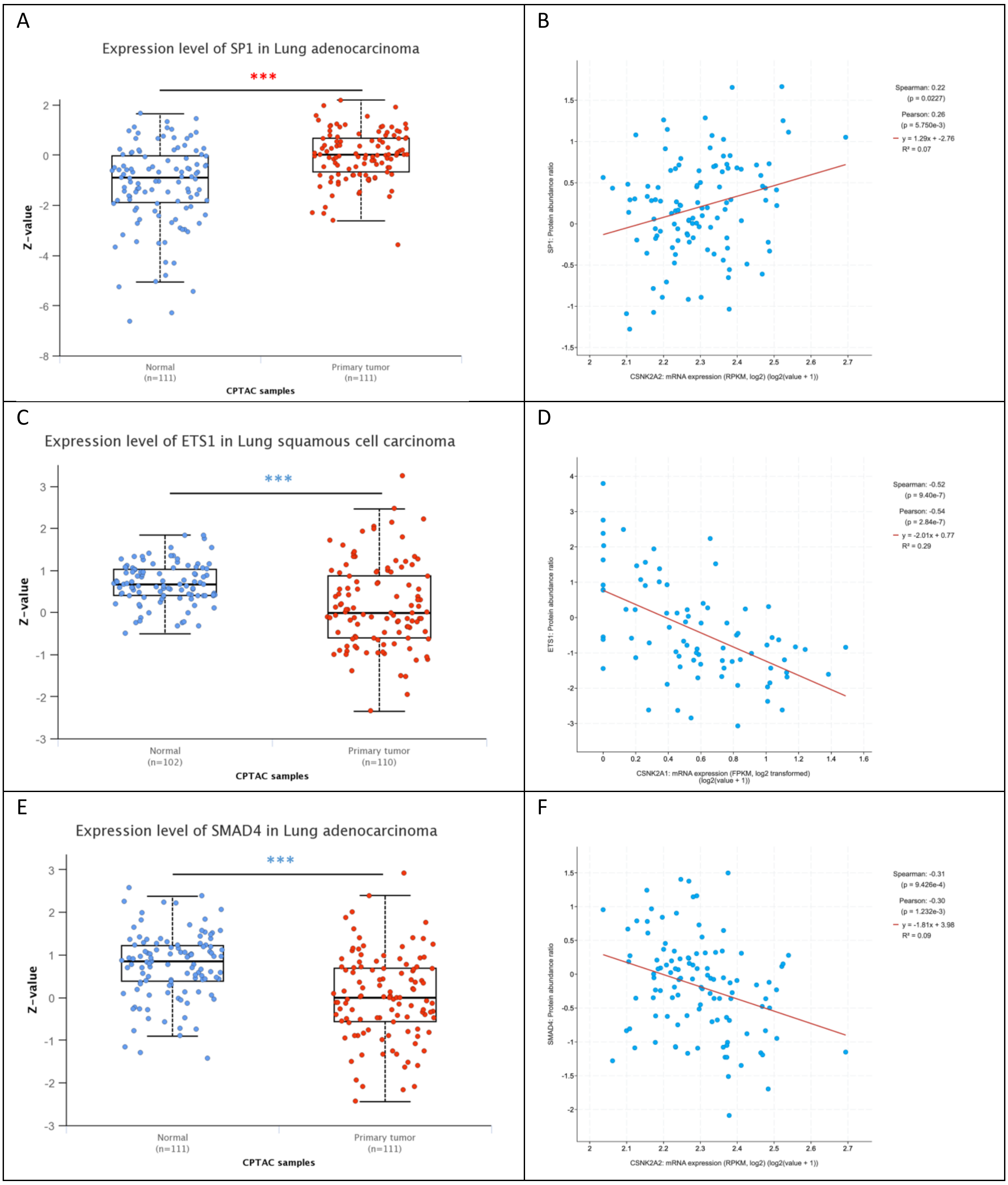

**SF.5 CK2 subunits expression across clinical cancer cohorts.** A) CSNK2A1 B) CSNK2A2 C) CSNK2B. The number of samples is represented below of each tumor subtypes. Groups with less than 15 samples are considerate non-significant for statistical comparison. Normal tissue (N) is highlighted in blue, while Tumor tissue (T) in red. Statistical significance is represented with red asterisks when the expression in tumor tissues is significative higher than normal tissues, and its representation in blue the opposite. TCGA tumor subtypes are described in the section Materials and methods.

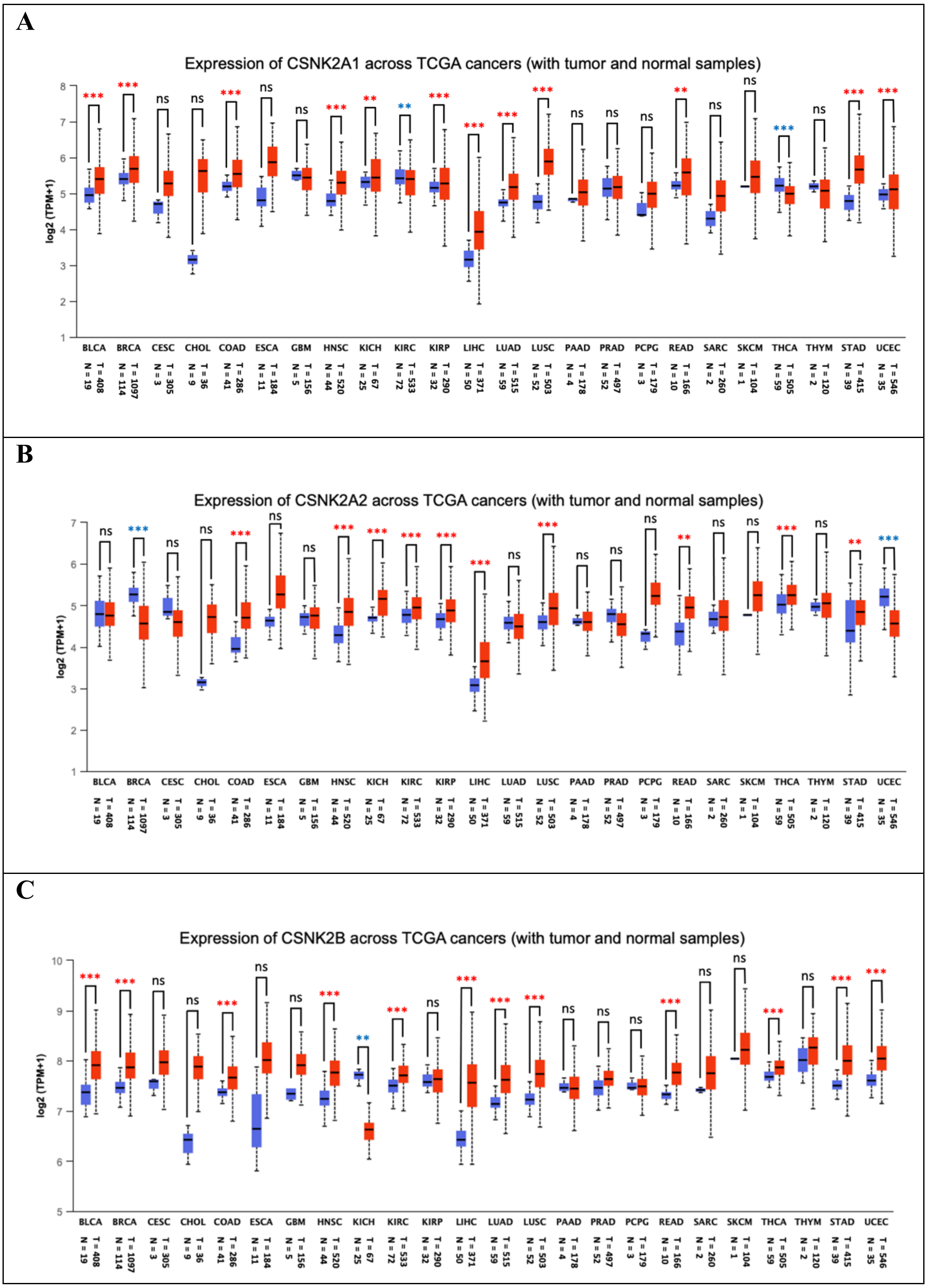

**SF.6 CSNK2 subunits gene expression across normal tissue, tumor and lung cancer metastasis.** Gene expression profile of A) CSNK2A1, B) CSNK2A2 and C) CSNK2B in normal (green), tumor (red) and metastatic (yellow) lung cancer samples; D) Expression density plot of CSNK2 genes in lung cancer, samples are represented as follow: normal (red), tumor (green) and metastatic (blue). Data analysis and representation obtained from TNMplot database.

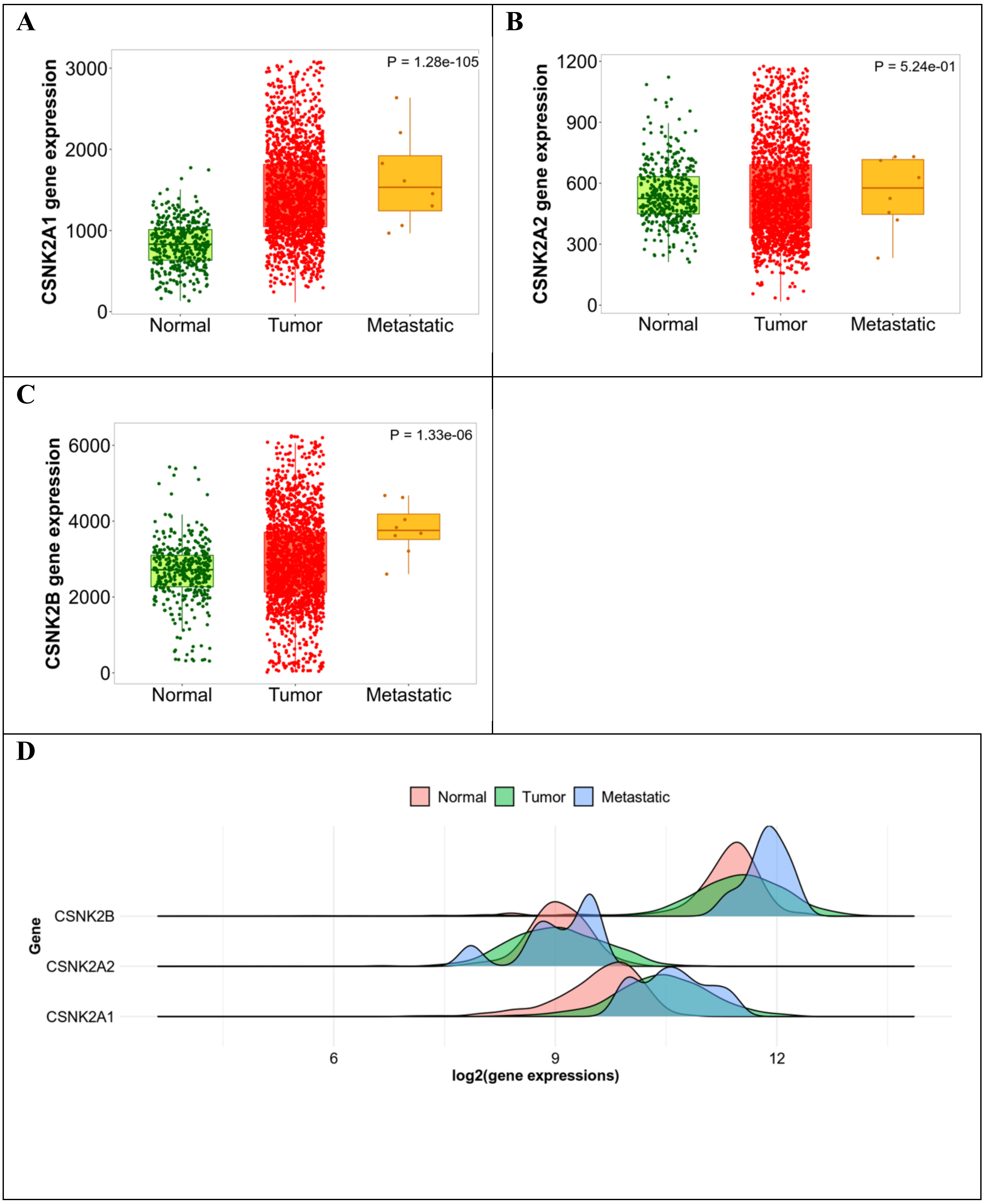

**SF.7 Meta-analysis of CNSK2A2 deregulation vs overall survival in 20 independent cohorts in NSCLC.** Global risk and heterogeneity scores are listed at left bottom of each chart. Curated studies in A) LUAD and B) LUSC.

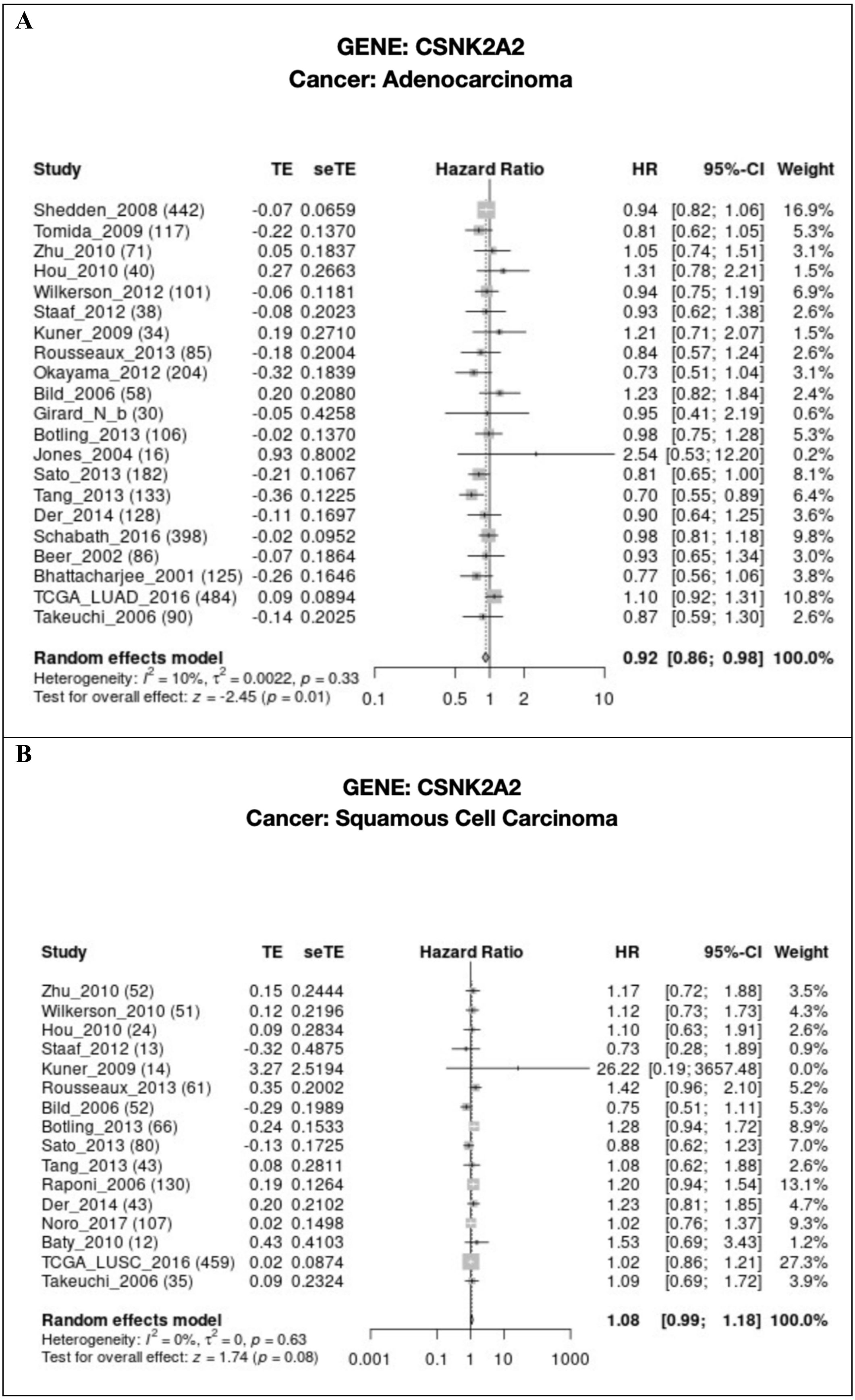

**SF.8 CSNK2A2 and CSNK2B over-expression correlates with selected immunosuppressive tumor infiltrating cells in different clinical cancer cohorts.** A) Correlation of CSNK2A2 expression and CAFs / MDSCs throughout different TCGA cancer subtypes; Detailed analysis of CAFs / MDSCs infiltration levels and CSNK2A2 expression in B) CAFs and C) MDSC samples. Each analysis was normalized according the purity of the samples. TCGA tumoral subtypes are described in the section Materials and methods.

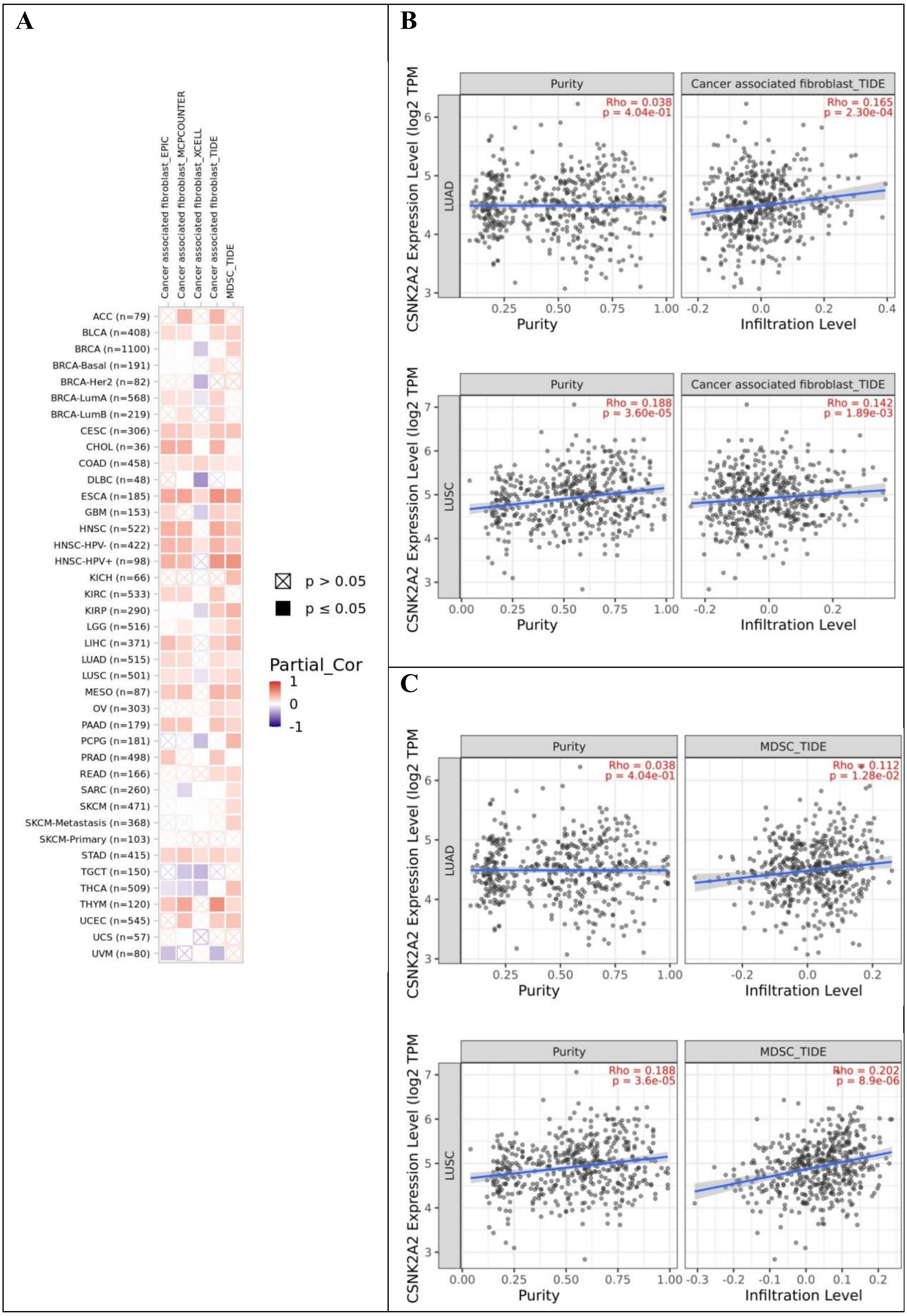

**SF.9 DepMap analysis of available pan-cancer cell lines for CK2 dependency scores.** Cell lines are represented as black dots. Negative values indicate that gene depletion affects viability/proliferation of each matched cell line. Red arrows from graphs are highlining lung tumoral tissue. Default Mix Ratio (CRISPR 60: shRNA 40) (C: R=60:40). DepMap for A) CSNK2A1, B) CSNK2A2 and C) CSNK2B. D). Substitute gene analysis for CSNK2B according efficacy/selectivity and functional similarity parameters. Red and orange dots are for CSNK2B and CSNK2A1 genes, respectively E) t-SNE plot for CSNK2B.

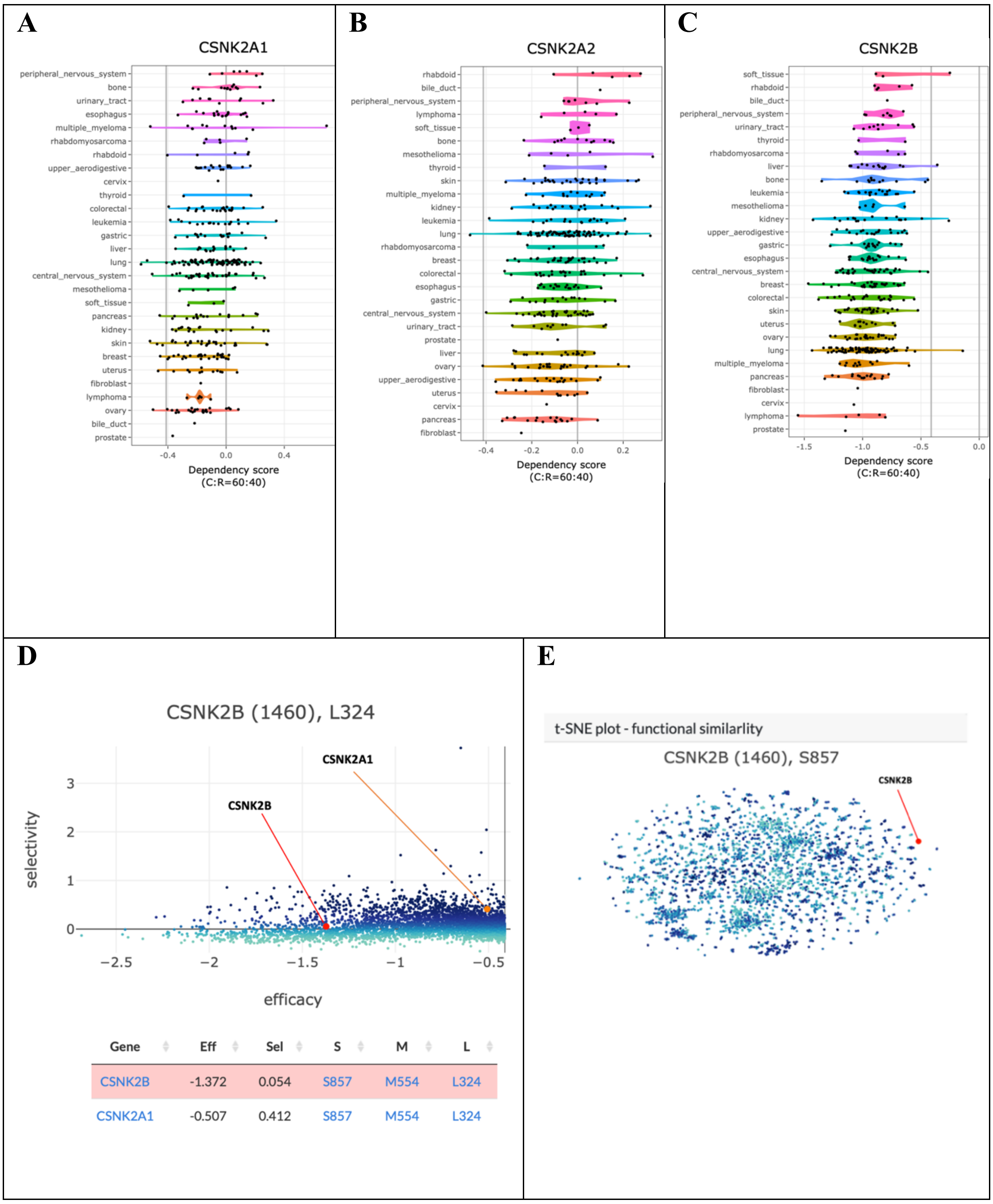

**SF.10 IHC detection of CK2 subunits and phosphorylation on selected CK2 substrates NPM1 in LUSC specimens and corresponding para-neoplastic tissues.** Representative results for each CK2 subunits are shown in tumor (A) or para-neoplastic tissues (B); C) Total NPM1 Protein and NPM1s125 phosphorylation in the same samples. Representative images were taken at 200X fold magnification.

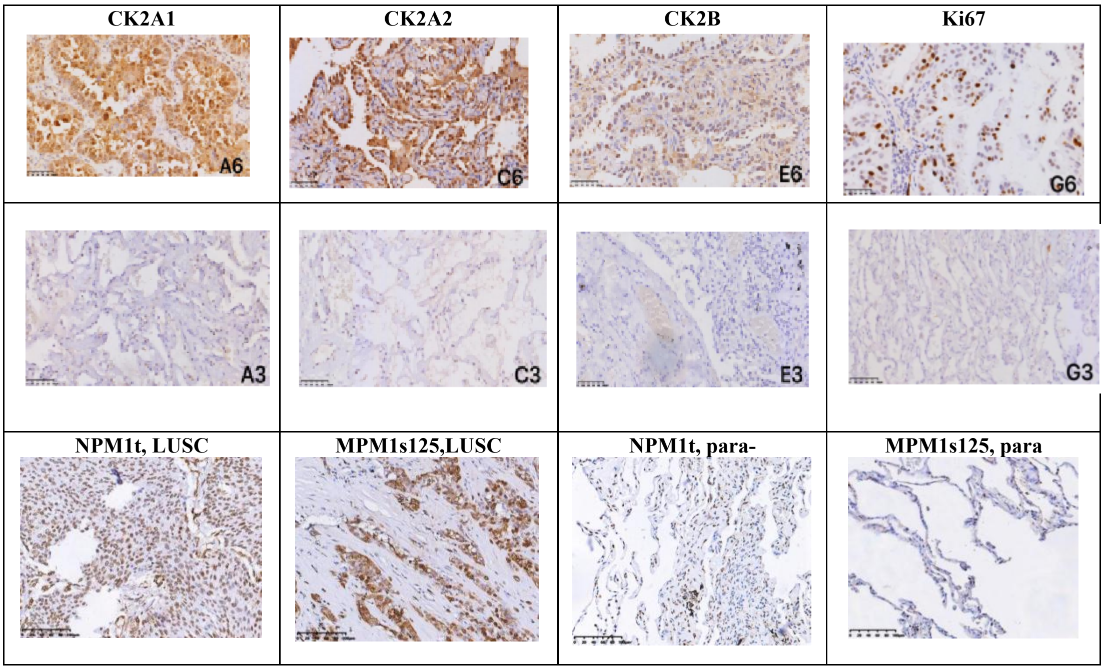

**ST.1 Correlation of relevant pathophysiological features with CSNK2A1 subunits levels in a NSCLC cohort containing LUAD and LUSC patient specimens**

| Pathophysiological features | Cases (n=134) | CK2A1 | | | | Positive rate(%) | Z | P |
|---|---|---|---|---|---|---|---|---|
| | | - | + | ++ | +++ | | | |
| Age | | | | | | | | |
| ≤60 | 64 | 14 | 26 | 12 | 12 | 78.2 | -0.495 | 0.621 |
| >60 | 70 | 16 | 24 | 15 | 15 | 77.2 | | |
| Gender | | | | | | | | |
| Male | 77 | 17 | 24 | 18 | 18 | 77.9 | -1.257 | 0.209 |
| Female | 57 | 13 | 26 | 9 | 9 | 77.2 | | |
| Smoking | | | | | | | | |
| Yes | 41 | 5 | 13 | 14 | 9 | 87.8 | -2.197 | 0.028* |
| No | 93 | 25 | 37 | 13 | 18 | 73.1 | | |
| Histology | | | | | | | | |
| LUAD | 103 | 29 | 44 | 17 | 13 | 71.8 | -4.894 | 0.000** |
| LSCC | 31 | 1 | 6 | 10 | 14 | 96.8 | | |
| Tumor size | | | | | | | | |
| ≤3cm | 110 | 27 | 47 | 19 | 17 | 75.5 | -3.733 | 0.000** |
| >3cm | 24 | 3 | 3 | 8 | 10 | 87.5 | | |
| Lymphatic metastasis | | | | | | | | |
| Yes | 18 | 4 | 4 | 5 | 5 | 77.8 | -0.979 | 0.321 |
| No | 116 | 26 | 46 | 22 | 22 | 77.6 | | |
| Pleural invasion | | | | | | | | |
| Yes | 12 | 3 | 3 | 1 | 5 | 75.0 | -0.828 | 0.407 |
| No | 122 | 27 | 47 | 26 | 22 | 77.9 | | |
| STAS | | | | | | | | |
| Yes | 44 | 12 | 17 | 7 | 8 | 72.7 | -1.059 | 0.289 |
| No | 90 | 18 | 33 | 20 | 19 | 80.0 | | |
| Differentiation | | | | | | | | |
| Low | 46 | 10 | 13 | 7 | 16 | 78.3 | -1.785 | 0.074 |
| High-medium | 88 | 20 | 37 | 20 | 11 | 77.3 | | |
| Clinical stage | | | | | | | | |
| I | 104 | 26 | 45 | 17 | 16 | 75.0 | -3.257 | 0.001** |
| II-IV | 30 | 4 | 5 | 10 | 11 | 86.7 | | |

***P*<0.05； ***P*<0.01**

**ST.2 Correlation of relevant pathophysiological features with CSNK2A2 subunits levels in a NSCLC cohort containing LUAD and LUSC patient specimens**

| Pathophysiological features | Cases (n=134) | CK2A2 | | | | Positive rate(%) | Z | P |
|---|---|---|---|---|---|---|---|---|
| | | - | + | ++ | +++ | | | |
| Age | | | | | | | | |
| ≤60 | 64 | 6 | 29 | 14 | 15 | 90.6 | -0.431 | 0.667 |
| >60 | 70 | 10 | 26 | 27 | 7 | 85.7 | | |
| Gender | | | | | | | | |
| Male | 77 | 11 | 30 | 26 | 10 | 85.7 | -0.746 | 0.455 |
| Female | 57 | 5 | 25 | 15 | 12 | 91.2 | | |
| Smoking | | | | | | | | |
| Yes | 41 | 5 | 12 | 16 | 8 | 87.8 | -1.341 | 0.180 |
| No | 93 | 11 | 43 | 25 | 14 | 88.2 | | |
| Histology | | | | | | | | |
| LUAD | 103 | 15 | 47 | 25 | 16 | 85.4 | -2.681 | 0.007** |
| LSCC | 31 | 1 | 8 | 16 | 6 | 96.8 | | |
| Tumor size | | | | | | | | |
| ≤3cm | 110 | 15 | 45 | 33 | 17 | 86.4 | -1.281 | 0.200 |
| >3cm | 24 | 1 | 10 | 8 | 5 | 95.8 | | |
| Lymphatic metastasis | | | | | | | | |
| Yes | 18 | 2 | 7 | 6 | 3 | 88.9 | -0.224 | 0.823 |
| No | 116 | 14 | 48 | 35 | 19 | 87.9 | | |
| Pleural invasion | | | | | | | | |
| Yes | 12 | 1 | 5 | 5 | 1 | 91.7 | -0.016 | 0.987 |
| No | 122 | 15 | 50 | 36 | 21 | 87.7 | | |
| STAS | | | | | | | | |
| Yes | 44 | 7 | 21 | 9 | 7 | 84.1 | -1.471 | 0.141 |
| No | 90 | 9 | 34 | 32 | 15 | 90.0 | | |
| Differentiation | | | | | | | | |
| Low | 46 | 6 | 15 | 19 | 6 | 87.0 | -0.470 | 0.638 |
| High-medium | 88 | 10 | 40 | 22 | 16 | 88.6 | | |
| Clinical stage | | | | | | | | |
| I | 104 | 13 | 45 | 29 | 17 | 87.5 | -0.913 | 0.361 |
| II-IV | 30 | 3 | 10 | 12 | 5 | 90.0 | | |

***P*<0.05； ***P*<0.01**

**ST.3 IHC-based detection and quantification (CIS) of NPM1/B23 phosphorylation as a proxy for *in vivo* CK2 enzymatic activity in LUAD (n=66).** A specific antibody recognizing the CK2 phosphoacceptor residue Ser125 on NPM1/B23 was used to detect substrate phosphorylation in LUAD specimens and corresponding para-neoplastic tissues. The data shows number and percent of samples displaying a certain CIS. A fisher exact test was used to detect statistical significance ($P < 0.01$)

| Groups | p-B23 | | | |
|---|---|---|---|---|
| | Negative | Weak positive | Positive | Strong positive |
| lung adenocarcinoma | 0（0.0） | 3（4.5） | 14（21.2） | 49（74.3） |
| adjacent | 29（43.9） | 7（10.6） | 21（31.8） | 9（13.7） |
| | $P < 0.01$# | | | |

#Pearson? 2 Test

**ST.4 IHC-based detection and quantification (CIS) of NPM1/B23 phosphorylation as a proxy for *in vivo* CK2 enzymatic activity in LUSC (n=16).** A specific antibody recognizing the CK2 phosphoacceptor residue Ser125 on NPM1/B23 was used to detect substrate phosphorylation in LUSC specimens and corresponding para-neoplastic tissues. The data shows number and percent of samples displaying a certain CIS. A fisher exact test was used to detect statistical significance ($P < 0.01$)

| Groups | p-B23 | | | |
|---|---|---|---|---|
| | Negative | Weak positive | Positive | Strong positive |
| lung squamous cell carcinoma | 0（0.0） | 1（6.3） | 4（25.0） | 11（68.7） |
| adjacent | 5（31.3） | 4（25.0） | 5（31.3） | 2（12.5） |
| | $P$=0.02* | | | |

*Fisher exact test

**ST.5 pNPM1 correlation analysis vs CK2 subunits in LUAD (upper table) and LUSC (lower table)**

| | p-B23 | CK2A1 | CK2A2 | CK2B |
|---|---|---|---|---|
| p-B23 | 1.000 | | | |
| CK2A1 | 0.177 | 1.000 | | |
| CK2A2 | 0.274* | 0.478** | 1.000 | |
| CK2B | 0.053 | 0.372** | 0.173 | 1.000 |

*$P < 0.05$ **$P < 0.01$

| | p-B23 | CK2A1 | CK2A2 | CK2B |
|---|---|---|---|---|
| p-B23 | 1.000 | | | |
| CK2A1 | 0.533* | 1.000 | | |
| CK2A2 | 0.597* | 0.683** | 1.000 | |
| CK2B | -0.019 | 0.249 | 0.430 | 1.000 |

*$P < 0.05$ **$P < 0.01$
